# Enhanced Brain-Heart Connectivity as a Precursor of Reduced State Anxiety After Therapeutic Virtual Reality Immersion

**DOI:** 10.1101/2024.11.28.625818

**Authors:** Idil Sezer, Paul Moreau, Mohamad El Sayed Hussein Jomaa, Valérie Godefroy, Bénédicte Batrancourt, Richard Lévy, Anton Filipchuk

## Abstract

State anxiety involves transient feelings of tension and nervousness in response to threats, which can escalate into anxiety disorders if persistent. Despite treatments, 30%-50% of individuals show limited improvement, and neurophysiological mechanisms of treatment responsiveness remain unclear, requiring the development of objective biomarkers.

In this study, we monitored multimodal electrophysiological parameters: heart rate variability (high-frequency, low-frequency, LF/HF ratio), EEG beta and alpha relative power, and brain-to-heart connectivity in participants with real-life state anxiety. Participants underwent a therapeutic intervention combining virtual-reality immersion, hypnotic script, and a breath control exercise. Real-life state anxiety was captured using the STAI-Y1 scale before and after the intervention.

We observed reduced anxiety immediately after the intervention in 16 out of 27 participants. While all participants, independently of their STAI-Y1 score, showed increased HRV low frequency power, only treatment-responders displayed increased overall autonomic tone (high and low frequency HRV), increased midline beta power and brain-to-heart connectivity. Notably, the LF/HF ratio showed a significant linear relationship with anxiety reduction, with higher ratios linked to greater therapeutic response.

These findings suggest that increased cognitive regulation of brain-to-heart connectivity could serve as a biomarker for therapeutic efficacy, with elevated midline beta power facilitating improved cardiac tone in responders.

**Significance Statement:** Elevated state anxiety can escalate into debilitating disorders, such as generalized anxiety disorder, yet treatment efficacy remains inconsistent, and reliable biomarkers predicting therapeutic outcomes are lacking. This study identifies key neural and physiological markers linked to effective anxiety reduction following a virtual reality-based non-pharmacological intervention in healthy participants with increased state anxiety. Anxiety reduction is associated with increased midline beta power, heightened heart rate variability (LF/HF ratio), and enhanced brain-to-heart connectivity. These findings highlight the role of brain-to-heart modulation and autonomic nervous system functioning in therapeutic response. By highlighting these biomarkers, this research aims at advancing our understanding of anxiety treatment mechanisms and offering insights into the development of biomarker-driven, scalable interventions.

## Introduction

Anxiety disorders, including generalized anxiety disorder (GAD), panic disorder, social anxiety disorder, and phobias, are among the most prevalent mental health conditions worldwide, affecting millions of individuals across the lifespan (1). Anxiety disorders affect more than 4% of the world population and their prevalence has increased by 55% in the last 20 years (1, 2). State anxiety, in contrast to trait anxiety related to personality traits, refers to a transient emotional state characterized by feelings of apprehension, tension, and nervousness in response to a perceived threat. Prolonged exposure to stressors, combined with maladaptive coping strategies, can result in persistent state anxiety, which may lead, over time, to chronic anxiety disorders (3). An escalating portion of the global population is experiencing heightened exposure to transient states of anxiety due to societal and environmental factors. This issue impacts various demographics, including but not limited to young active adults (4), students (5), urban dwellers (6), individuals living with chronic conditions (7) and informal caregivers of individuals with chronic conditions (8). The populations affected are significant and notably heterogeneous, rendering targeted solutions difficult. Typically, interventions only occur when stress levels escalate to clinically significant levels of anxiety and depression. Unfortunately, one-third of persons with anxiety disorders are unresponsive to pharmacological (9) or cognitive and behavioral therapy (CBT)-based treatments (10). Non-responders are individuals who do not exhibit clinically significant symptom improvements following treatment (11). The difficulties in treatment efficacy can be, at least in part, attributed to the lack of consideration for inter-individual variability as well as the multifaceted nature of anxiety (11, 12). While knowledge in neurobiological, genetic, and cellular markers of anxiety and treatment response has been gaining exponential traction (13, 14), the study of neural and cardiac biomarkers remains essential to provide real-time, non-invasive measures of bodily responses. These neural and cardiac biomarkers can be objectively monitored and may offer more immediate insights into the mechanisms underlying the efficacy of treatments.

A prominent theory posits that anxiety disorders arise from autonomic dysfunction (15). The autonomic nervous system (ANS) controls involuntary physiological functions, including heart rate. It consists of parasympathetic and sympathetic branches, respectively associated with physiological relaxation and arousal. The heart is controlled by both parasympathetic and sympathetic systems and *heart rate variability* (HRV), the variability between successive heartbeats plays an important role in mental as well as physiological health (16). High HRV is, in this sense, a protective feature, while low HRV has been associated with stress (15, 17). Individuals with Generalized Anxiety Disorder (GAD) often exhibit a relative deficit in vagally mediated HRV (18). Moreover, decreased HRV was observed in both non-anxious controls and patients during episodes of worry, suggesting that lower HRV can occur in response to stress, regardless of baseline anxiety levels (19). Anxiety disorders have been linked to increased cardiovascular morbidity and mortality (20). HRV has thus been exponentially used as a biomarker of anxiety disorders (21). The high-frequency (HF) component of HRV is associated with parasympathetic activity, while the low-frequency (LF) component reflects both sympathetic and parasympathetic activation and is not specific to sympathetic activation (22, 23). A meta-analysis investigating the relationship between stress and HRV spectral components summarized that decreased HRV during stress is likely associated with reduced parasympathetic activity, as indicated by a decline in HF spectral power (17). In summary, high HRV and elevated HF spectral power are indicative of a flexible and adaptive physiological system that supports homeostasis, while a rigid system (low HRV, low HF spectral power) exhibits greater difficulty in adapting to external stressors.

The ANS is regulated by the central autonomic network (CAN), a large-scale brain network. CAN involves the activation of medial cortices; notably the midcingulate cortex, subgenual and pregenual anterior cingulate cortices (sgACC, pgACC), extensively involved in emotional processes, precunueus that is related to self-referential processes (24), as well as the insula, amygdala, ventral tegmental area, and hypothalamus. The CAN regulates parasympathetic and sympathetic functions through medullary regions. These medullary regions include the dorsal motor nucleus of the vagus nerve, which controls parasympathetic activity, and the caudal ventrolateral medulla, responsible for sympathetic control, that is activated when external stressors challenge homeostasis (25). Dysfunction in the CAN, similarly to ANS dysfunction, has been linked to cardiac disease and anxiety disorders (26). The CAN plays a critical role in processing stressful stimuli, engaging higher-order cognitive regions involved in cognition, emotion, and autonomic regulation. EEG research has found a relationship between increased alpha band power and attention regulation (27). Parallelly, whole-brain increased beta band power was correlated to increased attentional control and top-down regulation after experimentally induced psychosocial stress (28). Additionally, frontal and midline beta power increase has been related to sustained attention (29).

Medial brain regions contain hubs of emotion regulation, such as the medial prefrontal cortex (mPFC) (30) as well motor and somatosensory regions. The somatosensory system, in particular, has gained recognition for its involvement in emotion regulation. For instance, increased beta neuromodulation via transcranial Alternating Current Stimulation (tACS) in the somatosensory region (S1)—part of the midline region—has been associated with positive emotional ratings (31). Moreover, distinct emotions, often described as “gut feelings” or “palpitations of the heart”, have been mapped to specific areas in the somatosensory homunculus (32), suggesting that motor areas contribute to more than just movement control. Beta bursts are also implicated in sensory processing, facilitating the coordination between sensory and motor functions (33).

Since state anxiety impacts both the autonomic and central nervous systems, studies of brain-heart interactions and brain-heart functional coupling have become valuable for understanding the functional manifestations of homeostatic variations (25, 34). Functional brain-heart coupling refers to the dynamic coordination of neural and cardiac activity, reflecting the interdependence of their functions. Recent models have investigated the directionality of brain to heart or heart to brain interactions using autoregressive methods (34). Its modulations have been associated with trait anxiety under a stressful situation (35). During a stressful task mimicking a preparation for an oral exam, authors evidenced a decrease in brain to heart connectivity in participants specifically during the stressful event (oral interrogation), at whole-brain level and for all investigated spectral bands (delta, theta, alpha and beta). In the context of emotional arousal, a prominent brain and heart connectivity study has shown that emotional reactions are initiated by ascending parasympathetic activation from the heart to the brain, with neural integration occurring through both ascending and descending interactions involving peripheral cardiac and central neural stimuli (36). In line with these ideas, the objectives of this study are twofold: 1) to elucidate the physiological and neural substrates associated with treatment responsiveness to the non-pharmacological intervention aimed at reducing state anxiety by analyzing data from all participants and contrasting the intervention effects with a control condition; and 2) to assess treatment response by comparing the effects of the virtual reality (VR) solution with a control condition within both responsive and unresponsive groups. This approach aims to identify potential biomarkers and elucidate mechanisms that may explain variability in responsiveness to the intervention. In the present study, we investigated the effects of a VR-based clinical solution designed to reduce state anxiety by immersing individuals in a *Zen Garden* virtual environment where natural landscapes are accompanied by a hypnotic script. The anxiolytic and analgesic effects of this therapeutic solution have been previously documented (37–39) although a fraction of patients remain non-responsive.

We explored peripheral and neural correlates of participants undergoing the therapeutic solution, as well as the directionality of brain-heart connectivity, by monitoring neurophysiological metrics. Cardiac (HF, LF, LF/HF ratio) and EEG (alpha and beta) correlates, along with brain-heart interactions, were investigated. We hypothesized that treatment-responsiveness would be related to the degree of brain-heart coupling, which would suggest improved flexibility and adaptability in maintaining homeostasis.

## Results

This study employed high-density EEG and high-resolution ECG to examine the effects of a VR-based therapeutic solution on state anxiety. Participants watched two videos with a VR headset in a randomized order. The therapeutic solution was a *Zen Garden* immersive environment, and the other a control neutral video showing American cities, both videos had a duration of approximately 20 minutes. The *Zen Garden* immersive environment was accompanied by a hypnotic script that guides the participant through the environment and begins with a breath control exercise (four seconds of inspiration and six seconds of expiration) until 188 seconds. The control neutral video presented aerial views of American cities, featuring relevant descriptions without emotional content.

Participants completed the short form of the State-Trait Anxiety Inventory (40) before and after viewing the Zen Garden video. A decrease in STAI-Y1 classified participants as ‘responsive,’ while unchanged or increased scores categorized them as ‘unresponsive’ (Supplementary Fig. 1). Baseline anxiety levels did not differ significantly between groups, indicating the absence of the pre-treatment stratification (Supplementary Fig. 2).

The ECG data collected during the study enabled the calculation of heart rate variability (HRV), whose variations are closely linked to stress and relaxation levels (17). We analyzed the spectral components of HRV, including low frequency (LF), high frequency (HF), and the LF/HF ratio, as these features provide insights into autonomic balance and emotional regulation (41).

Our EEG analyses focused on midline alpha (8-13 Hz) and beta (13-30 Hz) relative power, given their significance in reflecting state-dependent changes (42, 43) (Supplementary Fig. 3). Midline alpha activity is evidenced to be associated with relaxation (44) while midline beta activity relates to alertness and attentional engagement (28). By concentrating on these metrics, we can effectively explore the relationship between physiological relaxation/stress responses and attentional processes.

### Condition dependent autonomic, neural and psychometric responses in all participants

First, physiological metrics were compared between the *Zen Garden* condition and the control condition across all participants (n = 27, Fig. 1). As expected, STAI-Y1 scores were decreased after the therapeutic condition (mean ± SD = 27.7 ± 6.0 as opposed to 32.1 ± 10.4, p=0.023, Fig. 1, a), indicating a populational reduction in state anxiety. During the Zen Garden immersion, all data points (LF, HF, LF/HF ratio, midline beta and midline alpha relative band power) were averaged after excluding the initial paced breathing exercise epoch. This epoch was removed as voluntary control of breathing influences neural and cardiac activity (45).

**Figure 1.**
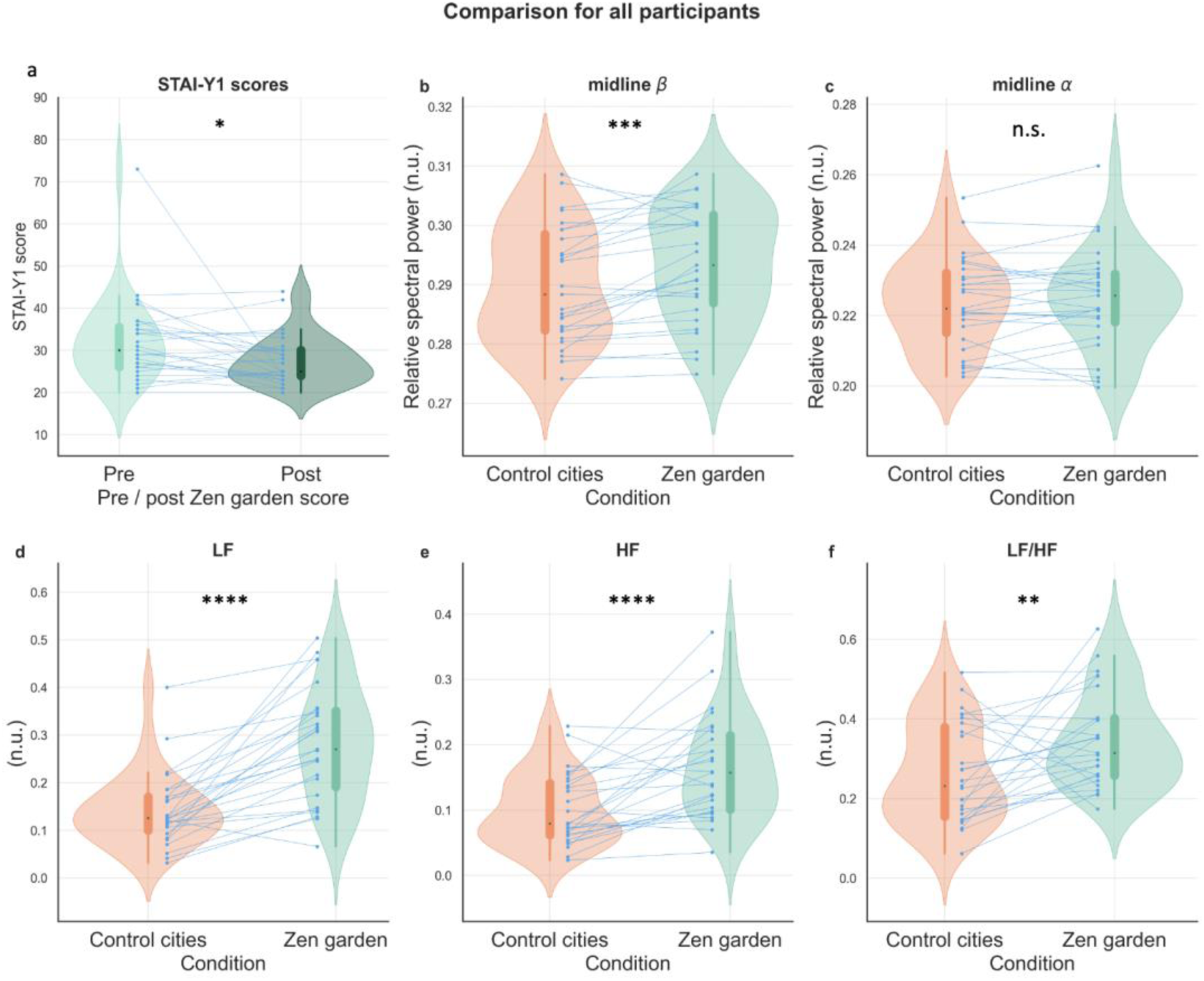
Conditions effects on STAI-Y1 ratings, neural and cardiac correlates. **a** STAI-Y1 scores of participants before and after immersion in the *Zen Garden* virtual environment (p=0.023, *W*=230, *r*=-0.44, 95% CI [−0.70, −0.11]). **b,c Relative band spectral power across conditions. b** Midline beta (p=0.0007, *W*=328, *r*=0.22, 95% CI [−0.09, 0.50]) **(c)** midline alpha (p=0.637). **d, e, f Cardiac measures differences across conditions**. **d** LF (p<0.0001, *W*=374, effect size *r*=0.70, 95% confidence interval (CI) [0.43, 0.85]), **e** HF (p<0.0001, *W*= 351, *r*=0.51, 95% CI [0.22, 0.72]), **f** LF/HF ratio (p=0.008, *W*=300, *r*=0.38, 95% CI [0.06, 0.63]). *Zen Garden* and control cities description conditions depicted in green and orange, respectively. Each participant’s data is represented by connecting blue points (N=27 participants). The boxplot inside the violin plot corresponds to the interquartile range, the median is depicted with a black dot, the vertical green and orange lines correspond to the probability density function. STAI-Y1 = State-Trait Anxiety Inventory, n.u. = normalized unit, LF = low frequency, HF = high frequency, * ≤ 0.05, ** ≤ 0.01, *** ≤ 0.001, **** ≤ 0.0001 p value with Wilcoxon signed rank test, adjusted with False Discovery Rate correction. STAI scores are commonly classified as ‘no or low anxiety’ (20–37), ‘moderate anxiety’ (38–44), and ‘high anxiety’ (45–80).

Regarding midline EEG correlates, relative beta (13-30 Hz) spectral power significantly increased in the therapeutic condition (mean ± SD = 0.294 ± 0.01 compared to 0.290 ± 0.01 in the control condition, p=0.0007, Fig 1, b), while no significant differences were observed in relative alpha (8-13 Hz) spectral power between conditions (mean ± SD = 0.225 ± 0.0143 compared to 0.224 0.0131 in the control condition, p=0.637, Fig. 1, c). ECG-derived metrics, LF, HF and LF/HF ratio, showed significant differences between conditions (Fig. 1, d,e,f). The condition of interest had a strong effect on mean LF values (mean ± SD = 0.281 ± 0.119 compared to 0.142 ± 0.078 in the control condition, p<0.0001, Fig 1, d), mean HF values (mean ± SD = 0.164 ± 0.078 compared to 0.101 ± 0.055 in the control condition, p<0.0001, Fig 1, e) and the LF/HF ratio (mean ± SD = 0.347 ± 0.120 compared to 0.263 ± 0.125 in the control condition, p=0.008, Fig 1, f). All these HRV components increased during the therapeutic condition, suggesting enhanced autonomic regulation and physiological relaxation during this phase.

### Group dependent differences in autonomic and neural response to the therapeutic condition: responsive and unresponsive participants

While the general condition effect was significant, it was primarily driven by the subgroup whose STAI-Y1 anxiety score substantially decreased (i.e. the responders) (Supplementary Fig. 1). To better understand these group-specific differences, the effects of the therapeutic condition were observed within each group (responders and non-responders), comparing *Zen Garden* mean values to control mean values in each subgroup (Table 1). LF of HRV showed significant condition differences for both responsive and unresponsive participants (Table 1). For responsive participants, differences were highly significant (Table 1, p=0.0003), and for unresponsive participants, differences were also marked (Table 1, p=0.01). Notably, only the group of responsive participants exhibited significantly higher mean HF values (Table 1, p=0.002) and an elevated averaged LF/HF ratio (Table 1, p=0.004) during the therapeutic condition compared to the control condition. In contrast, the non-responders showed no significant changes in either HF or LF/HF ratio between the therapeutic and control conditions (Table 1, HF: p=0.053, LF/HF ratio: p=0.641), although a trend toward significance was notable in HF. A distinct HF peak was observed during the controlled breathing phase, across participants (Fig. 2, a), with increased HF values throughout the *Zen Garden* condition for responsive participants (p=0.002, see Table 1 and Fig. 2, b,c), while declining to baseline levels in non-responders (p=0.053, see Table 1 and Fig. 2, d,e). A similar pattern emerged for the LF component of HRV, with a peak during paced breathing and maintained levels throughout the *Zen Garden* video for responsive participants, compared to a decrease following the peak for unresponsive participants (Supplementary Fig. 4). This dynamic was also observed for LF/HF ratio results (Fig. 3), with a distinctive peak in the removed breathing exercise epoch in all participants (Fig. 3, a). Responsive participants displayed increased LF/HF ratio value throughout the immersion in the therapeutic environment (Fig. 3, b), with statistically significant increase of averaged LF/HF values (p=0.004, see Table 1 and Fig. 3, c). Unresponsive participants, on the contrary, had a sharp decline of LF/HF ratio to control values after the breathing exercise (Fig. 3, d) and no difference across conditions (p=0.64, see Table 1 and Fig. 3, e). LF dynamics are illustrated in the Supplementary Figure 4.

**Figure 2.**
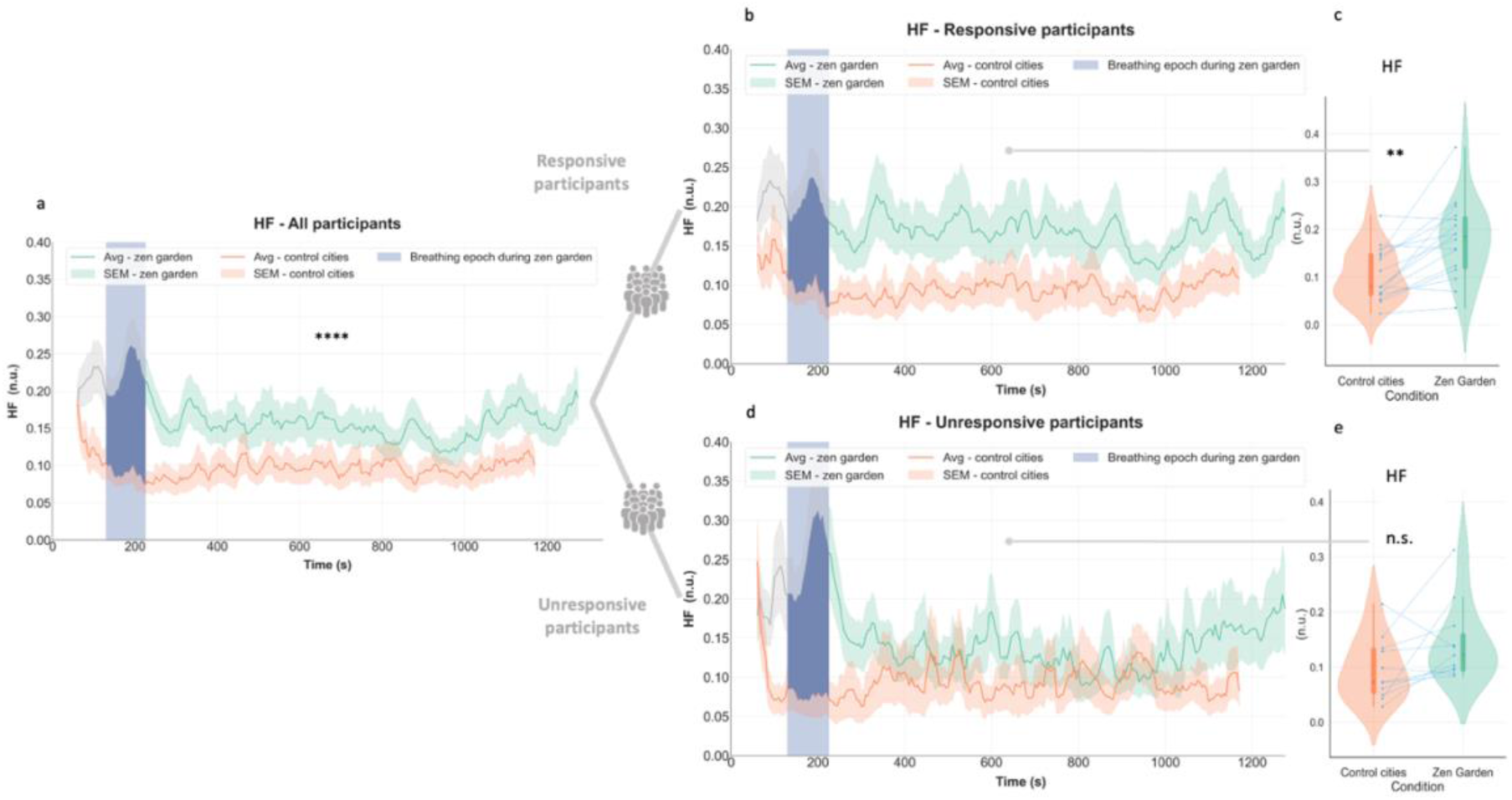
ECG HF spectral power response in each subgroup. **a** Temporal plots depict time-varying ECG HF differences across conditions in all participants (averaged HF spectral power is higher for *Zen Garden* condition, see Fig 1,e) **b** Responsive participants exhibited increase in HF spectral power values during the *Zen Garden* vs control conditions over 20-minute duration **c** Averaged HF spectral power increased during the *Zen Garden* condition (excluding the paced breathing induction epoch) compared the control video in the responsive group (p=0.002, *W*=130, *r*=0.58, 95% CI [0.17, 0.82]) **d** Unresponsive participants did not show difference in HF spectral power values during the *Zen Garden* compared to control condition over 20-minute duration **e** Unresponsive participants’ averaged HF spectral power did not significantly change during the *Zen Garden* condition compared the control condition (p=0.054). *Zen Garden* and control cities conditions depicted in green and orange, respectively. Temporal plots (b,d) depict datapoints for each window size=1 minute and step size=5 seconds, averaged ± standard error mean (SEM). Removed breathing exercise epoch is highlighted in blue. For (c,e) each participant’s data is represented by connecting blue points. The boxplot inside the violin plot corresponds to the interquartile range, the median is depicted with a black dot, the vertical green and orange lines correspond to the probability density function. N=27 all participants, n=16 responsive and n=11 unresponsive participants. Wilcoxon signed rank test, adjusted with False Discovery Rate correction.

**Figure 3.**
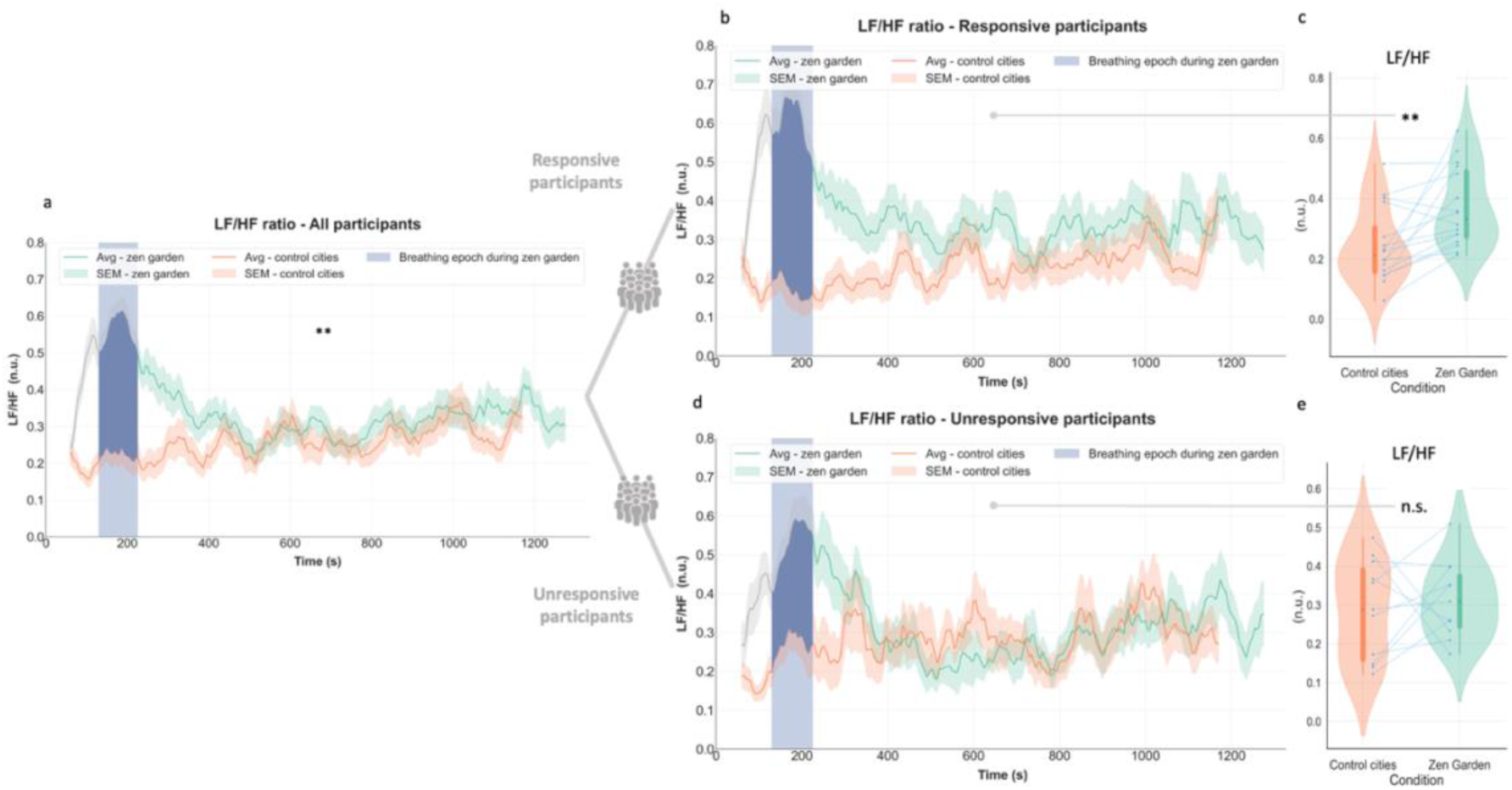
ECG HF/LF ratio response in each subgroup. **a** Temporal plots depict time-varying LF/HF ratio differences across conditions in all participants (averaged LF/HF ratio is higher for the *Zen Garden* condition, see Fig 1,f), **b** Responsive participants exhibited increase in LF/HF ratio values during the *Zen Garden* vs control conditions over 20-minute duration **c** Averaged LF/HF ratio increased during the *Zen Garden* condition (excluding the paced breathing induction epoch) compared the control video in the responsive group (p=0.004, *W*=126, *r*=0.57, 95% CI [0.16, 0.81]) **d** Unresponsive participants did not show difference in LF/HF ratio values during the *Zen Garden* compared to control condition over 20-minute duration **e** Unresponsive participants’ averaged LF/HF ratio did not significantly change during the *Zen Garden* condition compared the control condition (p=0.64). *Zen Garden* and control cities conditions depicted in green and orange, respectively. Temporal plots (b,d) depict datapoints for each window size=1 minute and step size=5 seconds, averaged ± SEM. Removed breathing exercise epoch is highlighted in blue. For (c,e) each participant’s data is represented by connecting blue points. The boxplot inside the violin plot corresponds to the interquartile range, the median is depicted with a black dot, the vertical green and orange lines correspond to the probability density function. N=27 all participants, n=16 responsive and n=11 unresponsive participants. Wilcoxon signed rank test, adjusted with False Discovery Rate correction.

**Table 1.** Comparison of each physiological metric across *Zen Garden* and control video conditions, within each sub-group.

| Physiological metric | Zen Garden and Control Cities comparison |  |  |  |
| --- | --- | --- | --- | --- |
|  | Responsive participants (n=16) |  | Unresponsive participants (n=11) |  |
|  | Zen Garden (m ± SD) | Control (m ± SD) | Zen Garden (m ± SD) | Control (m ± SD) |
| LF (n.u.) | **0.316 ± 0.123 \*\*\*** | **0.146 ± 0.084** | **0.229 ± 0.093 \*\*** | **0.135 ± 0.070** |
| HF (n.u.) | **0.178 ± 0.082 \*\*** | **0.103 ± 0.055** | 0.143 ± 0.070 | 0.096 ± 0.056 |
| LF/HF ratio | **0.370 ± 0.130 \*\*** | **0.244 ± 0.124** | 0.314 ± 0.099 | 0.289 ± 0.127 |
| EEG midline beta relative spectral power (n.u.) | **0.295 ± 0.009 \*\*** | **0.291 ± 0.009** | 0.291 ± 0.010 | 0.288 ± 0.010 |
| EEG midline alpha relative spectral power (n.u.) | 0.224 ± 0.014 | 0.223 ± 0.012 | 0.225 ± 0.015 | 0.224 ± 0.014 |

N=16 responsive participants and n=11 unresponsive participants. LF is significantly increased in the *Zen garden* condition for responsive participants ( $p=0.0003$ , $W=136$ , $r=0.74$ , 95% CI [0.38, 0.91]), as well as HF ( $p=0.002$ , $W=6$ , $r=0.58$ , 95% CI [0.17, 0.82]) and LF/HF ratio ( $p=0.004$ , $W=126$ , $r=0.57$ , 95% CI [0.16, 0.81]). On the contrary, for unresponsive participants, only the LF values are increased in the *Zen Garden* condition ( $p=0.01$ , $W=3$ , $r=0.59$ , 95% CI [0.06, 0.86]). Midline beta relative spectral power is increased during the *Zen Garden* condition in treatment-responders ( $p=0.009$ , $W=121$ , $r=0.250$ , 95% CI [-0.17, 0.59]). There is no difference across condition for midline alpha relative power ( $p=0.54$ in the responsive group and $p=0.96$ in the unresponsive group). Statistical analyses are performed with Wilcoxon signed rank test, adjusted with False Discovery Rate correction.

This breathing stimulus-driven dynamic was also visible in regard to midline beta EEG results (Fig. 4, a, all participants n=27). When observing each subgroup, only responsive participants had differences in beta band (13-30 Hz) relative spectral power across conditions (p=0.009, see Table 1 and Fig. 4, b,c), whereas unresponsive participants did not exhibit a statistically significant difference (p=0.118 see Table 1 and Fig. 4, e,f). Statistical results from other cortical regions are presented in Supplementary Figure 5 and Supplementary Figure 6. Subgroup analysis of the alpha band revealed that only responsive participants exhibited increased high alpha (10-13 Hz) relative power in occipital, temporal, parietal and central regions (Supplementary Fig. 7 and Fig. 8).

**Figure 4.**
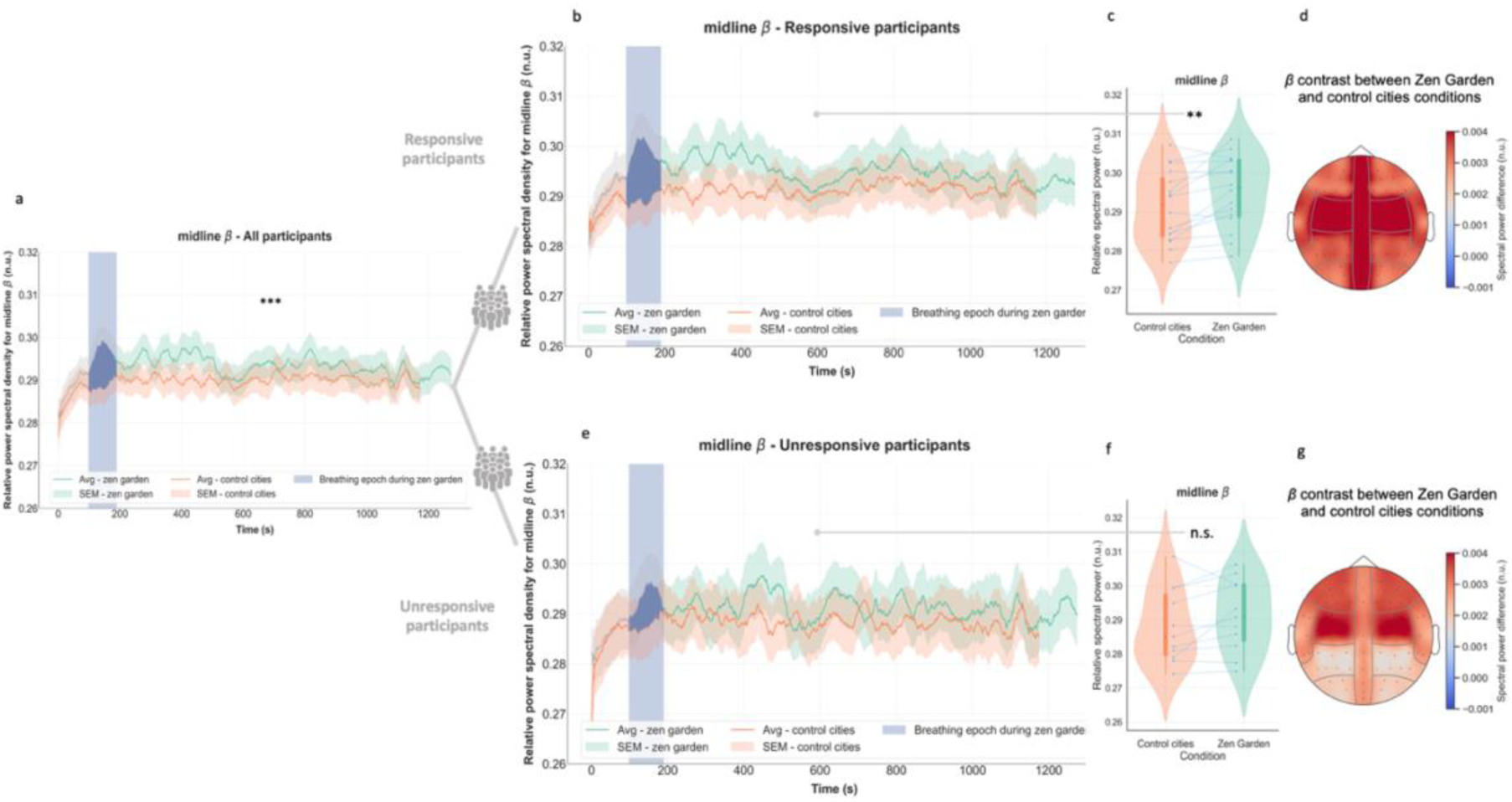
EEG beta band relative spectral power response in each subgroup. **a** Temporal plots depict time-varying midline relative beta differences across conditions in all participants (averaged midline beta relative spectral power is higher for *Zen Garden* condition, see Fig 1,b), **b** Responsive participants exhibited increase midline beta spectral power values during the *Zen Garden* vs control conditions over 20-minute duration **c** Averaged midline beta relative spectral power increased during the *Zen Garden* condition (excluding the paced breathing induction epoch) compared the control video in the responsive group (p=0.009, *W*=121, *r*= 0.250, 95% CI [−0.17, 0.59]) **d** Topographic representation of contrast between *Zen Garden* and control conditions in responsive group. **e** Unresponsive participants did not show difference in midline beta spectral power values during the *Zen Garden* vs control conditions over 20-minute duration **f** Unresponsive participants’ averaged midline beta relative spectral power did not significantly change during the *Zen Garden* condition (excluding the paced breathing induction epoch) compared the control condition (p=0.30) **g** Topographic representation of contrast between *Zen Garden* and control conditions in unresponsive group. For (d) and (g) groups of electrodes are averaged as in Supplementary Figure 3. *Zen Garden* and control cities conditions depicted in green and orange, respectively. Temporal plots (b,e) depict datapoints for each window size=1 minute and step size=5 seconds, averaged ± SEM. Removed breathing exercise epoch is highlighted in blue. For (c,f) each participant’s data is represented by connecting blue points. The boxplot inside the violin plot corresponds to the interquartile range, the median is depicted with a black dot, the vertical green and orange lines correspond to the probability density function. N=27 all participants, n=16 responsive and n=11 unresponsive participants. Wilcoxon signed rank test, adjusted with False Discovery Rate correction.

### Differences in brain-heart connectivity in responders and non-responders

Brain-heart coupling analyses were performed using the Sympatho-Vagal Synthetic Data Generation (SDG) model. Heart-to-brain (H2B) and brain-to-heart (B2H) coupling coefficients were calculated during the *Zen Garden* and the control video sessions for both subgroups (Fig. 5). This analysis focused on LF and HF HRV spectral power in relation to midline beta spectral power: HF2B, LF2B, B2LF, B2HF, averaged after the breathing exercise. For responsive participants, condition differences highlighted statistically significant increase from brain-to-heart HF (B2HF) (Fig. 5, a; p=0.002) and brain-to-heart LF coupling (B2LF) (Fig. 5, b; p=0.008) coefficients concomitant with decreases in heart-to-brain coupling coefficients: HF2B (Fig. 5, c; p<0.0001) and LF2B (Fig. 5, d; p<0.0001) in the *Zen Garden* condition, regardless of the heart frequency. These results in responsive participants strongly suggest an increased neural control of heart rate dynamics mediated by the midline beta frequency band, coupled with decreased control of neural midline beta activation by heart rate. In contrast, unresponsive participants exhibited a statistically significant decrease in the HF2B coefficient during the Zen Garden condition (Fig. 5, d; p=0.004) and did not show any condition differences in B2HF, B2LF or LF2B (Fig. 5 a,b,e). This difference may indicate that the condition effect is primarily mediated by brain-to-heart directionality.

**Figure 5.**
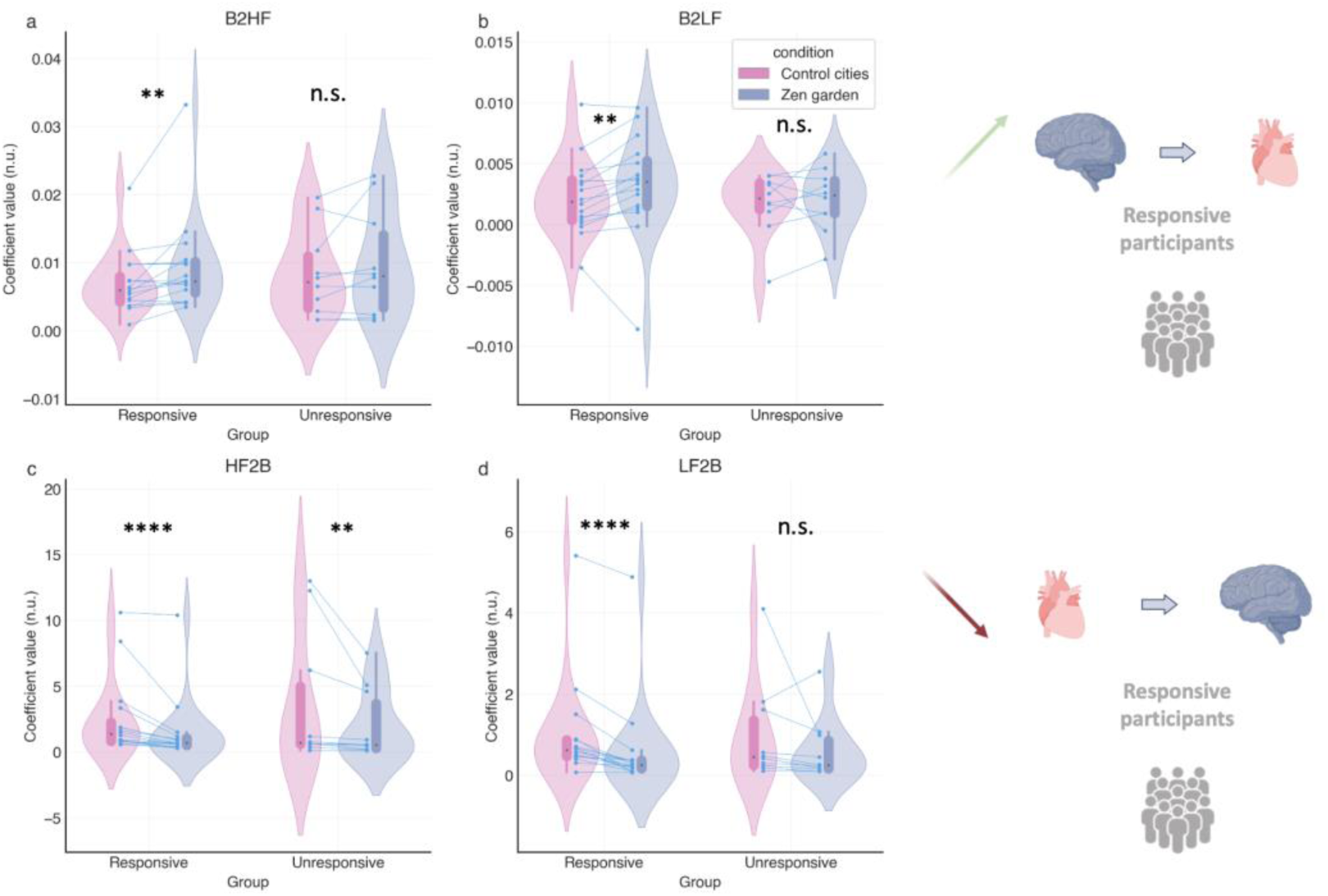
Brain-to-heart coupling coefficients across conditions in responsive and unresponsive groups. **a** Brain to high-frequency spectral power (B2HF) in each group across conditions; B2HF increases in the responsive group (p=0.002, *W*=10, *r*=0.27, 95% CI [−0.152, 0.603]) and there is no difference in the non-responsive group (p=0.49). **b** Brain to low-frequency (B2LF) coupling coefficient values in each group across conditions; increased B2LF coefficient values (p=0.008, *W*=18, 95% CI [−0.156, 0.605]) during the *Zen Garden* condition in the responsive groups; no changes in the unresponsive group (p= 0.49). Summary graphical representation of main brain to heart connectivity findings in responsive participants. **c** High-frequency spectral power to brain (HF2B) is significantly reduced in the responsive participants (p<0.0001, *W*=136, *r*=-0.47, 95% CI [−0.74, −0.05]) as well as in the unresponsive group (p= 0.008, *W*=55, *r*=-0.28, 95% CI [−0.66, 0.22]) in the *Zen Garden* condition. **d** Low-frequency to brain (LF2B) coupling coefficient values are significantly reduced in the responsive group (p<0.0001, *W*=136, *r*=-0.47, −0.59, 95% CI [−0.84, −0.15]) in the *Zen Garden* condition. No significant difference was observed for the unresponsive group (p=0.13). Summary graphical representation of main heart to brain connectivity findings in responsive participants. *Zen Garden* and control cities description conditions depicted in pink and purple, respectively. Each participant’s data is represented by connecting blue points. The boxplot inside the violin plot corresponds to the interquartile range, the median is depicted with a black dot, the vertical pink and purple lines correspond to the probability density function. Diagonal top green arrow corresponds to an increase in connectivity, diagonal bottom red arrow indicates to a decrease in connectivity. N=16 responsive participants, n=10 unresponsive participants. Wilcoxon signed-rank test, adjusted with False Discovery Rate correction. Graphical depiction created in https://BioRender.com.

### LF/HF ratio-based prediction of magnitude of relaxation response

Statistical analyses were performed to detect the relationship between individuals’ physiological responses and the magnitude of differences in STAI-Y1 scores before and after the intervention (Fig. 6). 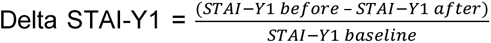 values were estimated using a linear regression model, with LF values, HF values, and LF/HF ratio as predictors. Among these, only the LF/HF ratio yielded statistically significant predictions (p=0.008). The positive estimate indicates that for each unit increase in the LF/HF ratio, the magnitude of response (calculated as the difference, or delta, in anxiety score divided by the baseline value) is expected to increase. A separate model was constructed to predict response magnitude (delta STAI-Y1) based on the midline beta and the midline alpha relative spectral power values. Estimations were not statistically significant. The third model, which investigated the influence of brain-heart coupling coefficients on anxiolytic responses, did not yield any statistically significant regressors either.

**Figure 6.**
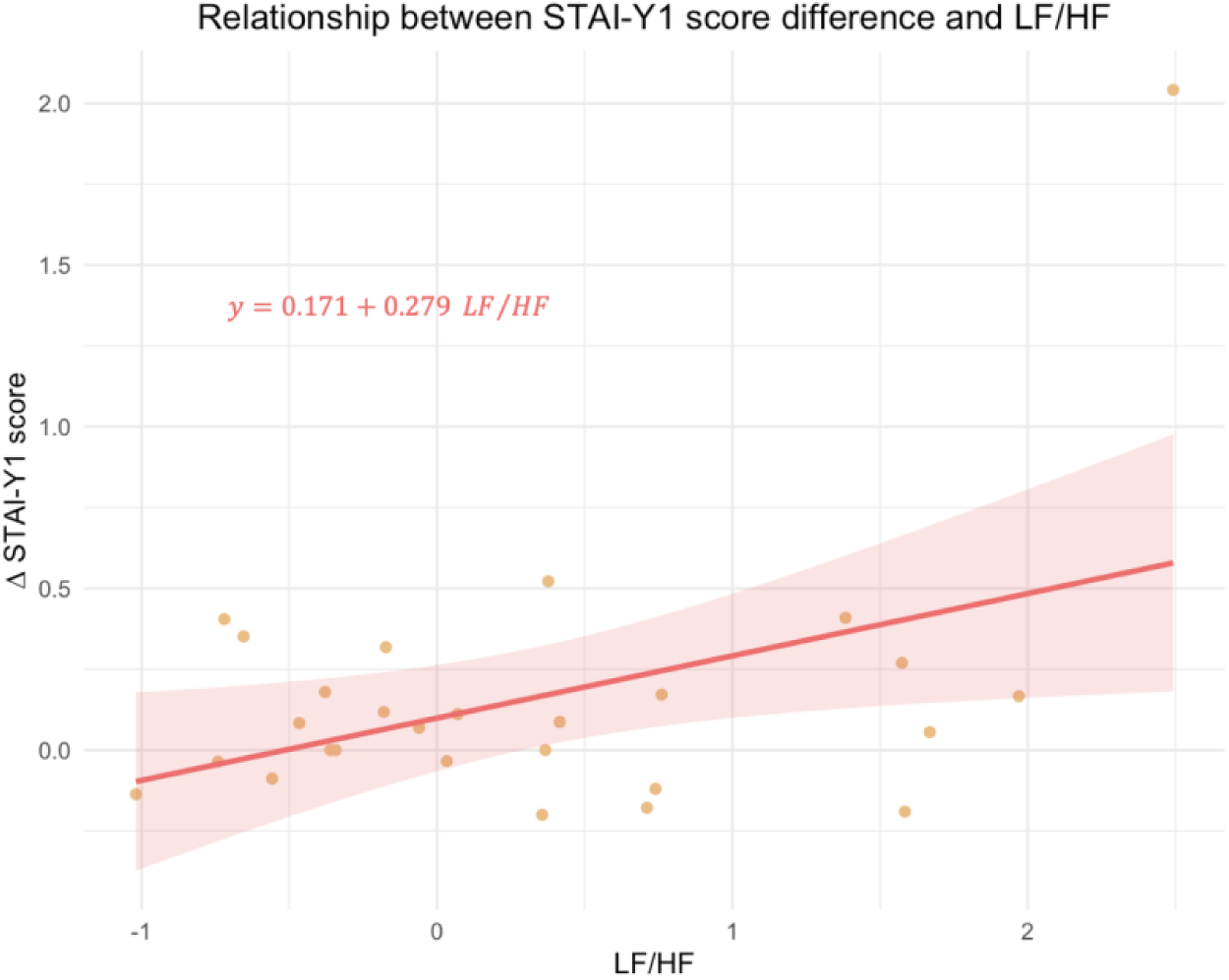
STAI-Y1 response magnitude in relation to LF/HF ratio. Delta STAI-Y1 is calculated as 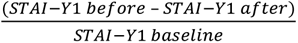. Each point represents an individual participant’s LF/HF ratio (n.u.) and corresponding delta STAI-Y1 score. LF/HF values yielded statistical significance to STAI-Y1 difference (p=0.008, *t*= 2.90, *Cohen’s d*=0.615, 95% CI [0.18, 1.05]). The regression line, in red, represents the best linear fit for the data (the model’s linear coefficient for LF/HF is 0.279). The shaded region around the regression line indicates the standard error of the estimate. N=27 participants.

## Discussion

In this present study we explored the neural and autonomic mechanisms associated with state anxiety reduction following a non-pharmacological intervention. The first objective was to pinpoint physiological responses to a VR-based solution aiming at reducing state anxiety, assessed with the State-Trait Anxiety Inventory. The second objective was to differentiate the physiological correlates between responsive and unresponsive participants. As expected, a fraction of participants was non-responsive to treatment with unchanged state anxiety scores after the therapeutic intervention. While the treatment effect is strongly associated with cardiac features, and all participants exhibited increased low frequency (LF) HRV band power during the relaxing therapeutic VR video, only the parasympathetic high frequency (HF) HRV spectral power and midline beta EEG values discriminated the therapeutic response in individuals classified as responsive. Responsive participants, identified as those who displayed a decrease in state anxiety scores, exhibited increases in the HF power, the LF/HF ratio, and midline beta relative band power. Notably, at the interindividual level, the increase in the LF/HF ratio was significantly related to the intensity of anxiety reduction. In terms of brain and heart interactions, responsive participants displayed enhanced directional brain-to-heart connectivity, with increased midline beta EEG activity linked to enhanced LF and HF cardiac power, while unresponsive participants did not display this connectivity (Fig. 7).

**Figure 7.**
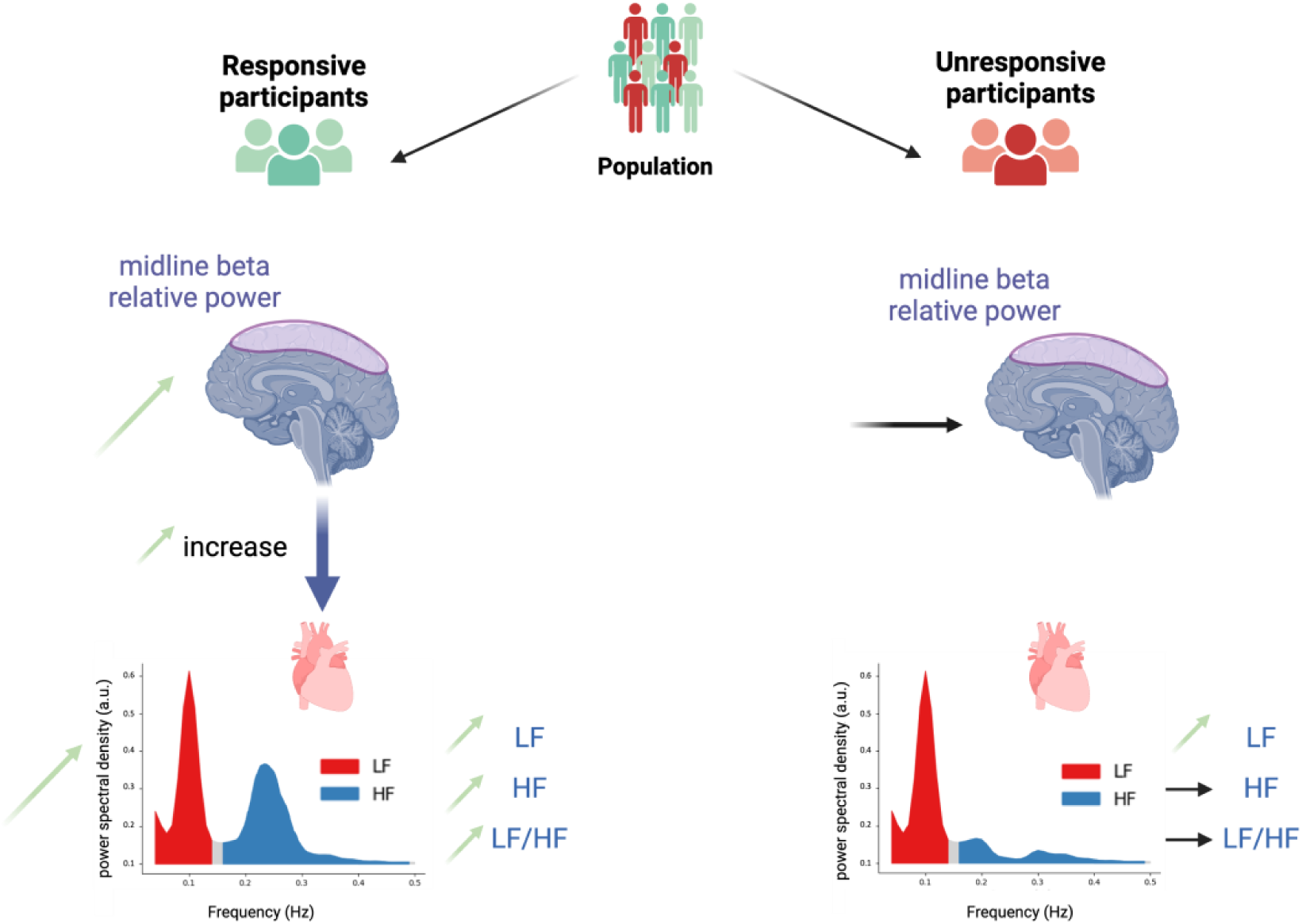
Mechanistic explanation for treatment response-related group differences. Responsive participants exhibit increased midline beta relative spectral power connectivity with both the LF and the HF heart spectral powers. They display elevated midline beta relative spectral power, as well as increased LF, HF values and increased LF/HF ratio. In contrast, unresponsive participants show no significant changes in midline beta brain-to-heart connectivity, regardless of the HF or LF spectral power. They display no alterations in the midline beta relative spectral power, the HF spectral power, or the LF/HF ratio. The only statistically significant changes observed in this group are an increase in the LF heart spectral power and a decrease in the HF toward the midline beta spectral power (the latter not illustrated for clarity). Green arrows indicate a statistically significant increase in the *Zen Garden* condition compared to the control, while horizontal black arrows denote unchanged differences across conditions. Figure created with BioRender.com (Figure license: Sezer, I. (2024) https://BioRender.com/r10p104).

The state anxiety levels reported by participants were naturally occurring rather than experimentally induced, which underscores the real-world applicability of the intervention beyond laboratory settings. By integrating psychometric ratings, cardiac metrics, EEG data, and brain-heart connectivity, our multimodal study provides a comprehensive understanding of the intervention’s mechanism of action in participants classified as responsive to the therapeutic intervention. The convergence of these diverse physiological and neural measures offers a comprehensive and quantitative insight into how the treatment impacts responsive individuals.

The experimental paradigm of this study seeks to explain the physiological precursors of action of the non-pharmacological intervention. However, such an intervention produces a synergistic therapeutic effect, where the impact could arise from the combined influence of the hypnotic script, the natural setting, the calming sounds, and the virtual immersion. To disentangle these factors, future studies should investigate whether one or more of these components are driving the main effects.

### Role of ANS in anxiety regulation

Autonomic function plays a pivotal role in emotional regulation, particularly in relation to the stress response and the development of complex adaptive behaviors, as posited by prominent cardiac-centric theories such as the neurovisceral integration theory and the polyvagal theory (19, 46). Both theories underscore the importance of parasympathetic regulation in facilitating emotional resilience and behavioral flexibility.

Previous research has emphasized parasympathetic dominance, reflected by HF activation, as a key indicator of relaxation (47, 48). However, our findings provide a more nuanced explanation of these cardiac markers, as we observed simultaneous increases in LF, HF, and in the LF/HF ratio in treatment responders. Notably, respiratory control plays a significant role in the global increase in cardiac spectral oscillations, as 10-second respiratory cycles align with a 0.1 Hz resonant frequency peak within the LF band. This synchronization at 0.1 Hz, termed cardiac coherence, reflects the coupling between the cardiac and respiratory systems. This synchronization contributes to self-regulation and improves emotional regulation, stress management and cognitive function. As a result, this phenomenon promotes overall psychological and physical well-being (49, 50). The discriminant effect of the treatment in our study appears to be mediated by concurrent increases in HF and the LF/HF ratio, observed exclusively in the responder group. This suggests that the breath control exercise alone does not fully account for the observed effects. Instead, the observed outcomes are likely tied to the participants’ ability to engage top-down regulatory cognitive processes that activate this optimal autonomic state throughout the therapeutic session.

Of note, although all cardiac metrics increased in responders, the LF/HF ratio was the only significantly predictive metric of the intensity of the anxiolytic response. Previous studies coined LF/HF ratio as a straightforward marker of sympathetic/parasympathetic balance (51). Our results challenge this assumption. While unresponsive participants displayed increases exclusively in the LF band; suggesting a sympathetic activation in line with their lack of relaxation; the increased LF/HF ratio in responders does not imply a shift toward sympathetic dominance. Instead, the interpretation of LF/HF requires caution, as both LF and HF are influenced by a complex interplay of autonomic mechanisms, with LF incorporating both sympathetic and parasympathetic inputs. Recent literature has raised concerns regarding the over-reliance on LF/HF ratio as a straightforward measure of sympathovagal balance, advocating for a more nuanced understanding of its physiological underpinnings (52). Given HRV’s importance in assessing physiological and emotional states of individuals, as well as its potential as a transversal diagnostic marker (21, 53), it becomes paramount to develop new methods that could elucidate the complex and intricate dynamics of autonomic subsystems.

### Contribution of non-pharmacological factors into CNS and ANS interaction

The therapeutic intervention contains several facets that could explain the observed anxiolytic response. A recent systematic review (54) highlights that immersion in VR environments has been strongly associated with relaxation and increased HRV, mediated by the autonomic nervous system. In this line, our findings support the efficacy of non-pharmacological interventions, which aligns with previous research. For instance, a study on older adults demonstrated a significant reduction in anxiety scores after immersive VR reminiscence therapy, with participants reporting decreased anxiety levels after each session (55). Additionally, another pilot study found that participants engaging in a mindfulness-based VR intervention reported notable reductions in state anxiety compared to a control group (56).

The beneficial effects of naturalistic environments on mental health have also been supported by a meta-analysis linking exposure to natural landscapes, as opposed to urban environments, with positive outcomes in emotion regulation and well-being (57). Another study examined the physiological effects of exposure to natural versus urban videos, revealing that participants displayed increased sustained attention and enhanced stress recovery during the viewing of natural environments (58). That particular study suggested that even viewing natural videos can serve as a proxy for being outdoors, highlighting the potential benefits of virtual exposure to nature for mental health, especially for individuals unable to go outside, such as those undergoing clinical interventions.

Practices like hypnosis and meditation, despite their differences, share common neural responses linked to heightened executive control, attention regulation, and altered states of consciousness (59). These practices often employ breath control to help regulate attention. In the immersive experience within the context of this study, participants are shielded from external distractions, which aids their focus on visual and auditory stimuli happening in the immersive environment, mirroring mindful attention practices. Although the sustained differential state marked by prolonged increase in beta markers evidenced during the relaxing therapeutic environment may not perfectly align with traditional hypnotic or meditative correlates, all these practices involve heightened attention control facilitated by reduced external stimuli. Meditative practices have been evidenced to increase HRV (60) and cardiac-neural interactions (61).

### Midline beta activation and enhanced top-down brain-to-heart regulation

Midline beta activation has been shown to mediate therapeutic responses, particularly by enhancing cognitive control (29) and sustaining attention regulation (28). This control aids in the integration of internal and external stimuli, as mental processes are influenced by visceral interoceptive feedback (41). The differential response between individuals may reflect variations in cognitive and autonomic integration, with participants exhibiting greater flexibility more effectively engaging attentional processes in response to interventions (62).

In the context of brain-heart interaction, responsive participants’ metrics revealed elevated midline beta relative spectral connectivity towards HF and LF heart rate oscillations. In this responsive group, these were paired with decreased ascending LF and HF heart rate spectral power connectivity towards midline beta relative spectral power. In contrast, unresponsive participants did not show increased top-down connectivity. Taking these findings into account, we hypothesize that the response mechanisms operate via midline cortical control of the heart, leading to parasympathetic activation and increased autonomic tone (Fig. 7). The amygdala, a key node in the central autonomic network (CAN), is additionally involved in emotion regulation (63), while the hypothalamus, also part of the CAN contains both sympathetic and parasympathetic pre-ganglionic neuronal population, is likely involved in the observed vagal activation (26). In parallel, detailed alpha band analysis demonstrated significant increase in the high alpha relative spectral power in occipital, parietal, central and temporal regions, in line with decreased anxiety (64).

Our findings additionally align with previous brain-heart functional interplay literature stating that a stressful episode was associated with decrease in brain-to-heart connectivity (whole-brain and delta, theta, alpha and beta bands) (35). It is therefore plausible that, in contrast, increased brain-to-heart connectivity, as observed in our study, is associated with reduced state anxiety.

Midline regions, notably the medial prefrontal cortex (mPFC), part of the Default Mode Network (DMN), play a pivotal role in autonomic modulation, as demonstrated by a meta-analysis, linking midline activity to increased vagal tone and heart rate variability (HRV) (15). This enhanced vagal tone is crucial for inhibiting stress responses and improving emotional regulation (19). This neural system has been proposed to operate as a “super-system,” integrating perceptual, motor, interoceptive, and memory functions to form adaptive responses to gestalt representations of situations. This medial hub, integrating a wide range of information, ranging from primitive to higher-order neural processes, helps explain how increased midline beta findings are in line with multiple functions: sustained attention control, emotion regulation, sensory-motor integration, overall promoting well-being.

### Limitations and perspectives

This study is limited by its relatively small sample size (n=27), which may affect the generalizability of the findings. However, the highly significant results obtained from this sample underscore the substantial effect size of the treatment. Additionally, the classification of participants into responsive and unresponsive groups was based on post-hoc analyses, which may introduce bias. While the use of linear regression to predict the response magnitude addressed this issue, future studies should consider larger sample sizes and pre-defined criteria for participant classification, potentially leveraging the presently discovered physiological biomarkers of responsiveness. This treatment uses synergistic factors, strongly suggesting a significant effect from their combined utilization. As the therapeutic effect of the intervention was assessed in previous literature, the goal of the study was not to dissect each element but to understand the mechanism of action of the whole intervention. In line with this, future studies should assess the contribution of each of the factors (breath control exercise, hypnotic script, naturalistic environment immersion, mindful awareness strategies) to tailor the therapeutic intervention to each individual.

Despite these limitations, this research provides valuable insights into the effects of a therapeutic video in immersive VR, integrating multiple physiological measures with psychological self-reported ratings. This multimodal approach enhances our understanding of the intervention’s impact on everyday-life anxiety, a condition that is increasingly prevalent. The absence of a stress-inducing experimental paradigm strengthens the relevance of our findings, suggesting that the intervention may lower real-life stress levels and serve as a preventive measure for anxiety disorders.

### Conclusions

Midline beta and autonomic substrates show promising potential as biomarkers for predicting and monitoring treatment response. Specifically, the LF/HF ratio emerged as a key indicator of response magnitude, with higher ratios predicting greater reductions in state anxiety. Future research should further investigate the role of cardiac autonomic markers and the role of midline beta in facilitating the brain-heart communication essential for therapeutic efficacy, particularly in non-pharmacological interventions targeting elevated stress. Additionally, exploring individual differences in baseline autonomic function and neural connectivity may provide insights for personalizing treatments and reducing non-responsiveness (65).

Future research should also explore other underlying mechanisms driving the differential responses to therapeutic videos, examining genetic, psychological, or contextual factors that influence responsiveness. Investigating other types of meditative or nature-based stimuli could help identify specific features that maximize therapeutic benefits. Longitudinal studies on the long-term effects of regular exposure to these interventions on physiological and psychological health would be valuable. In the short term, the discovery of these physiological biomarkers, including sustained increases in HF and midline relative beta spectral powers, as well as elevated LF/HF ratio predictive of response intensity, could guide he development of neurofeedback and biofeedback-based therapeutic systems.

In conclusion, the therapeutic non-pharmacological intervention induced significant neural, cardiac, and neuropsychological changes, highlighting key biomarkers in participants classified as responsive. The findings support the potential of nature-based and meditative interventions in reducing anxiety by promoting autonomic balance. Personalized approaches should be considered to optimize the efficacy of such interventions, acknowledging that individual differences play a crucial role in therapeutic outcomes.

## Materials and Methods

### Participants

27 healthy volunteers (age range 24-59; mean age (±SD) = 37.30 ± 11.27; 14 women; 2 left-handed, 1 ambidextrous; mean years of tertiary education (±SD) 4.07 ± 2.23) were recruited via an online advertisement on the RISC platform (Réseau d’Information en Sciences de la Cognition) in France. Participants had no history of psychiatric or neurological conditions. Seven participants were excluded, resulting in an attrition rate of 21%. One participant was excluded due to a history of hypertension affecting electrocardiogram (ECG) signal quality, one for not completing the state anxiety questionnaire, and five participants were excluded because of uncorrectable high frequency ECG artifacts. Exclusions ensured data quality but may limit generalizability to those with similar ECG challenges. The study was approved by the Ethics Committee of Paris Sorbonne University (registration number ‘CER-2022-063’ approved by ‘Comité d’Ethique de la Recherche Sorbonne Université’). All participants gave informed consent and their personal data were pseudo-anonymized in compliance with ethical guidelines. Sample size determination, based on a review (66) of 297 HRV effect sizes (R; pwr package; Cohen’s d = 0.6; Alpha error probability: 0.05; power = 0.8), estimated that 24 participants would provide 80% power to detect a medium effect size (d = 0.6) in heart rate variability as an indicator of cardiac autonomic control in a paired design study.

### Experimental design

Participants watched 2 videos using a VR headset: Pico Neo 3 VR goggles and Bose QuietComfort 35 II headphones. The VR headset, connected to a PC via USB, displayed the videos through custom-made C# software (*NeuroRelaxMarkersV7*), which synchronized triggers sent to an Arduino microcontroller linked to the EEG recording system. The latency between VR events and triggers was under 100ms, meeting accuracy requirements. The therapeutic solution was a *Zen Garden* immersive environment (Healthy Mind, Paris, FRANCE), and the other condition consisted of a control neutral video showing American cities. Both videos had a duration of approximately 20 minutes (Supplementary Fig. 9). The *Zen Garden* immersive environment was accompanied by a hypnotic script (see Supplementary Table 1). The audio guided participants through the environment (Supplementary Fig.10), encouraging focus on visual elements and relaxation through mental simulations (e.g., the refreshing sensation of water droplets on the face). The beginning of the recording starts with a breath control exercise (4 seconds of inspiration and 6 seconds of expiration) until 188 seconds. The control neutral video showed several American cities with relevant descriptions of them. State anxiety was assessed using the 20-item STAI-Y1 (40), administered three times: before the protocol, between videos, and after the videos.

### Data acquisition

Electroencephalogram (EEG) data were acquired using a 64-channel BrainAmp system (Brain Products GmbH, Germany) with two stacked 32-channel units, following the 10-10 electrode placement (see Supplementary Fig. 3). Signals were recorded differentially to ground (Fpz) and referenced to FCz. Data were sampled at 1 kHz and band-pass filtered (0.016–80 Hz). ECG was recorded in parallel using Neuroline 720 Ambu electrodes (Ambu, France) at 1 kHz, with placements at the right clavicle (−1), left pelvis (+1), and upper back (ground). The experiment was conducted in a dark, soundproofed, and electromagnetically shielded environment within a Faraday cage to prevent external interference.

### ECG data processing

ECG inter-beat intervals (IBIs) were filtered using an EEGLAB Butterworth filter, between 0.5 and 30 Hz. R peaks from the QRS waves were detected using Pan-Tompkins’ toolbox (65, MATLAB), followed by visual inspection and automated correction of misdetections or ectopic beats via Neurokit2 (66, Python). The low-frequency (LF, 0.04–0.15 Hz) and high-frequency (HF, 0.15–0.4 Hz) components of HRV reflect mixed sympathetic/parasympathetic and parasympathetic nervous system activity, respectively. These components were computed using the heartbeat analysis section of the Synthetic Data Generation (SDG) model, as outlined by (69). To account for slow variations in HRV dynamics, these LF and HF calculations were performed using a sliding window of 1 minute and a step size of 5 seconds. The initial epoch of the *Zen Garden* condition, including the breathing exercise (<188 s), was excluded. To account for interindividual differences, each participant’s data were normalized using min-max scaling.

### EEG data processing

Ocular and movement artifacts were mitigated using wavelet denoising (pywt, Python). Each EEG channel was processed individually by decomposing the signal into wavelet coefficients using the Coiflet-3 (coif3) wavelet. Channel-specific thresholds were applied to the coefficients to suppress noise while preserving relevant signal components. After thresholding, inverse wavelet reconstruction was performed to obtain the denoised signal. EEG data were filtered (1–48 Hz) with a 50 Hz notch filter to remove powerline interference. Each channel’s denoised output and the removed artifacts were reconstructed into the original data structure using MNE (68, MNE-Python). Midline channels were selected (AFz, Fz, Cz, CPz, Pz, POz, Oz, Iz) as they are physiologically relevant for state-dependent analyses (see Supplementary Fig. 5, Fig. 6, Fig. 7, Fig. 8 for other regions’ results). EEG bands were extracted using a Finite Impulse Response (FIR) filter (MNE-Python) with a Hamming window: alpha (8–13 Hz), beta (13–30 Hz), theta (4–8 Hz), and low gamma (30–46 Hz). Spectral power was computed via Short-Time Fourier Transform (STFT, MATLAB) using a 1-second window with 50% overlap. Relative alpha and beta power in midline electrodes was calculated by normalizing each electrode’s spectral power to the total power of all four bands. Midline relative beta and alpha spectral frequency powers were averaged across electrodes. The initial epoch of the *Zen Garden* condition, including the breathing exercise (<188 seconds), was excluded, the remaining values were averaged for each participant.

### Brain-heart coupling calculation

Time series of beta power were extracted from midline EEG electrodes (AFz, Fz, Cz, CPz, Pz, POz, Oz, Iz), and time series of low-frequency (LF: 0.04–0.15 Hz) and high-frequency (HF: 0.15– 0.4 Hz) spectral power were derived from heart rate variability (HRV) using a sliding window approach (15-second window, 1-second step). To compute the directionality and strength of brain-heart coupling, coupling coefficients were estimated using the Synthetic Data Generation (SDG) model, as outlined by (69). The SDG model applies autoregressive transfer functions to evaluate time-varying, bidirectional connectivity between EEG spectral power and HRV components. For each window of 15 seconds, the SDG model estimated four coupling coefficients: HF2B (High Frequency to Beta) and LF2B (Low Frequency to Beta), calculated using autoregressive exogenous (ARX) models (MATLAB) where HRV components served as predictors of beta power dynamics. B2HF (Beta to High Frequency) and B2LF (Beta to Low Frequency), were calculated by normalizing EEG beta power and regressing HRV dynamics (LF and HF) against EEG-derived time-domain predictors generated using synthetic datasets. For each condition, the coefficients were averaged across all time points for each participant. The initial epoch of the *Zen Garden* condition (<188 seconds) was excluded, the remaining values were averaged. One non-responder participant was removed due to an HF2B outlier value (5– 95% range criterion). Their exclusion did not affect results.

### Statistical analyses and grouping

To assess the effect of conditions, first, all metric values were averaged for each participant in each condition, with Zen Garden means calculated after the controlled breathing phase (188 seconds after the start). Data across participants were standardized to z-scores. Since determining that data were not normally distributed, two-tailed Wilcoxon signed-rank tests were used to compare condition means of the following metrics: LF, HF, LF/HF, alpha and beta relative power. Multiple comparisons were corrected using the False-Discovery Rate (FDR) correction following the Benjamini-Hochberg procedure. STAI-Y1 scores before and after the *Zen Garden* video were compared using the Wilcoxon signed-rank test after confirming non-normal distribution. Participants with decreased state anxiety scores were classified as ‘responsive’ (n = 16; mean age = 36.31 ± 11.78; 9 women), while those with unchanged or increased anxiety scores were classified as ‘unresponsive’ (n = 11; mean age = 39.18 ± 12.13; 5 women). There were no statistical differences between the groups in age, sex, years of education, handedness, or the order in which videos were watched. Within each group, conditions mean differences for all metrics (LF, HF, LF/HF, alpha and beta relative band spectral powers) were compared using Wilcoxon signed-rank tests after testing for normality. Group comparisons were examined by comparing metrics (LF, HF, LF/HF, alpha and beta relative midline band power) within each group across *Zen Garden* and control conditions. Multiple comparisons were corrected using FDR correction. Test statistics (*W*), Cliff’s delta effect sizes, and confidence intervals for the effect sizes were reported (effsize package, R).

### Response magnitude calculation and linear modeling

State anxiety scores were calculated as 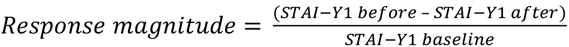, where before and after refer to pre and post-intervention measures. STAI-Y1 baseline was the score obtained prior to viewing both video stimuli. For participants who viewed the *Zen Garden* video first, baseline and pre-intervention scores were the same. Linear regression models were fitted to examine the relationship between heart physiological metrics (LF, HF, LF/HF ratio) and the magnitude of the anxiety response. Test statistics, Cohen’s d effect size, and the corresponding confidence intervals for the effect size were computed. A separate linear regression model evaluated the influence of relative midline alpha and midline beta spectral power on the calculated response magnitude. A third linear regression model evaluated the relation of brain heart coupling coefficients (B2HF, B2LF, HF2B, LF2B) to response magnitude.

## Acknowledgments

Authors thank D. Candia-Rivera (Paris Brain Institute) for their support in fitting the SDG model to the dataset and S. Gobbi for her support for customized VR software development.

## Data and code availability

All data required to reproduce our findings are publicly available at https://github.com/idilsezer/brain-heart_state-anxiety-biomarkers/tree/main/data and https://zenodo.org/records/14640287. The Python code for data processing and figure creation, along with R scripts for statistical analyses, are available at https://github.com/idilsezer/brain-heart_state-anxiety-biomarkers.

## Author Contributions

Conceptualization A.F.; Experimental investigation: A.F. I.S. P.M.; Data preprocessing: I.S. P.M. M.J.; Data analyses: I.S.; Visualizations: P.M. I.S. M.J. A.F.; Writing, original draft: I.S. A.F.; Writing review and editing: A.F. B.B. R.L. V.G. M.J.

## Competing Interest Statement

The therapeutic intervention and equipment used in this study include products developed by Healthy Mind, a private company with which Idil Sezer, Mohamad El Sayed Hussein Jomaa and Anton Filipchuk hold full or partial affiliation. Employees of Healthy Mind participated in the study design (A.F.) and data analysis (I.S., M.J.). This affiliation and the company’s involvement have been fully disclosed to all authors and participants. All findings reported in this study were collected, analyzed, and interpreted with strict adherence to scientific rigor to ensure objectivity and minimize potential bias associated with this affiliation. Importantly, this study was not intended to assess the efficacy of the therapeutic intervention (previously published) but to investigate the physiological mechanisms underlying its effects.

## Notes

https://github.com/idilsezer/brain-heart_state-anxiety-biomarkers.git

### Summary of Updates

The manuscript has been shortened. Figure 8 has been moved to supplemental.

