## Supplementary information for "Enhanced Brain-Heart Connectivity as a Precursor of Reduced State Anxiety After Therapeutic Virtual Reality Immersion"

**This PDF file includes:**

Figures S1 to S10
Table S1

**Other supporting materials for this manuscript include the following:**

Datasets S1 to S3, S5 to S8, datasets Figure 1 to 6, datasets Table 1

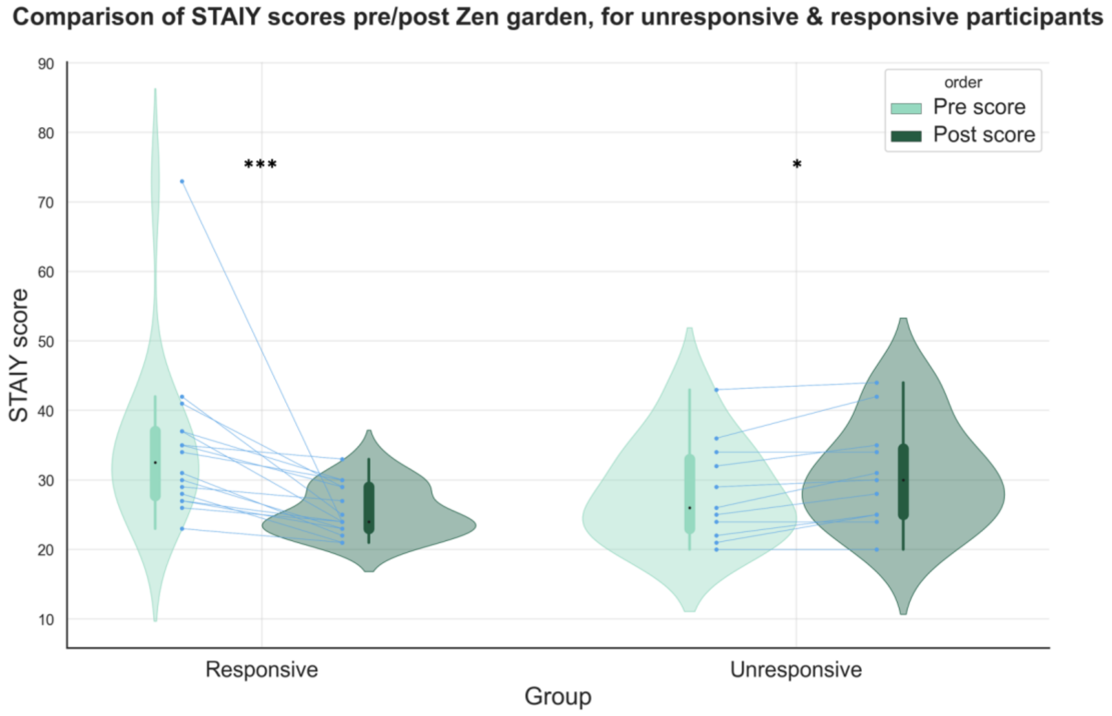


**Fig. S2. Zen Garden condition effects on psychometric ratings in each group.** STAI-Y1 state anxiety scores of responsive (p=0.0005, *W*=136, *r*=0.87, 95% CI [0.87, 0.89]) and unresponsive (p=0.01, *W*=0, *r*=-0.74, 95% CI [-0.88, -0.57]) participant groups. Control cities and *Zen Garden* conditions depicted in bright green and dark green, respectively. Each participant's data is represented by connecting blue points (N=16 responsive and n=11 unresponsive participants). The boxplot inside the violin plot corresponds to the interquartile range, the median is depicted with a black dot, the vertical bright and dark green lines correspond to the probability density function. Statistical analyses performed using Wilcoxon signed rank test.

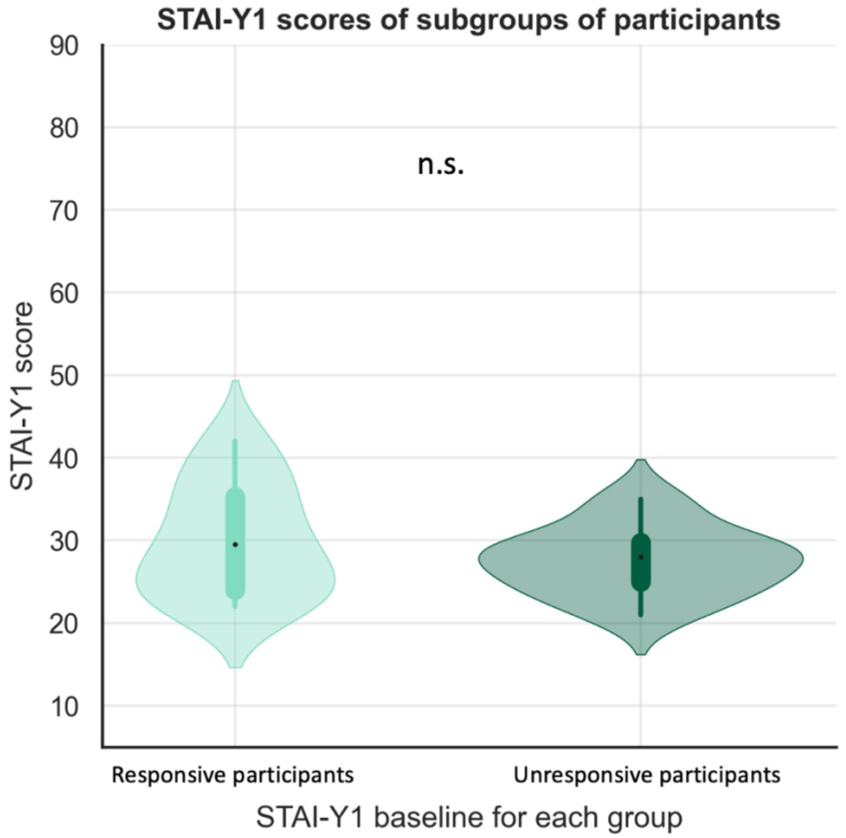


**Fig. S2. Baseline state anxiety levels.** Baseline STAI-Y1 state anxiety scores in participants classified as responsive or unresponsive (p= 0.71), before viewing *Zen Garden* or control videos. N=16 responsive and n=11 unresponsive participants. The boxplot inside the violin plot corresponds to the interquartile range, the median is depicted with a black dot, the vertical bright and dark green lines correspond to the probability density function. Statistical analysis performed using Mann Whitney U test.

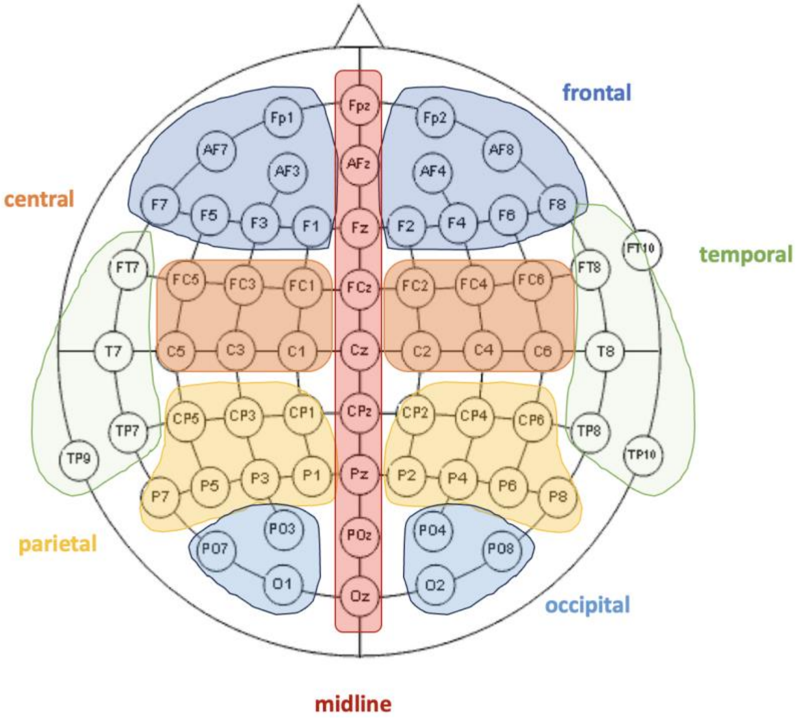


**Fig. S3. Grouping of electrodes by region for analyses.**
Frontal region electrodes: Fp2, AF4, AF8, F8, F6, F4, F2, Fp1, AF3, AF7, F7, F5, F3, F1
Temporal region electrodes: FT8, FT10, T8, TP8, TP10, FT7, FT9, T7, TP7, TP9
Central region electrodes: FC2, FC4, FC6, C6, C4, C2, FC1, FC3, FC5, C5, C3, C1
Parietal region electrodes: CP2, CP4, CP6, P6, P4, P2, CP1, CP3, CP5, P5, P3, P1
Occipital region electrodes: P8, PO8, PO4, O2, P7, PO7, PO3, O1
**Midline region electrodes: AFz, Fz, Cz, CPz, Pz, POz, Oz, Iz**

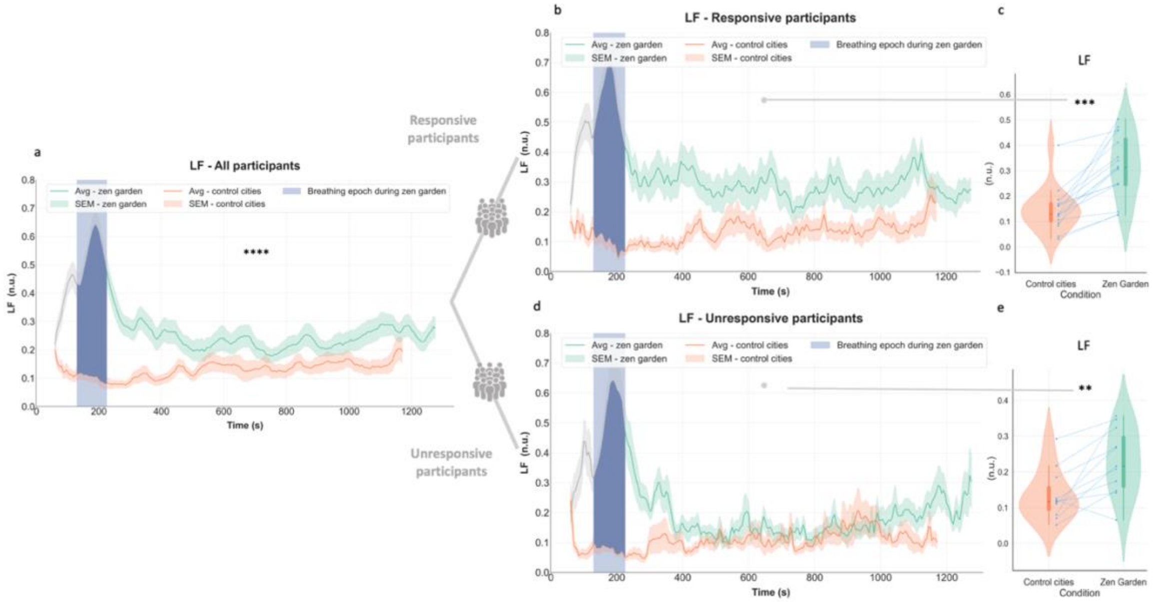


**Fig. S4. ECG LF spectral power response in each subgroup**. **a** Temporal plots depict time-varying ECG LF differences across conditions in all participants (see Fig 1, d for statistics) **b** Responsive participants exhibited increase in LF spectral power values during the *Zen Garden* vs control conditions over 20-minute duration **c** Averaged LF spectral power increased during the *Zen Garden* condition (excluding the paced breathing induction epoch) compared the control video in the responsive group (p=0.0003, *W*=136, *r*=0.74, 95% CI [0.38, 0.91]) **d** Unresponsive participants exhibited increase in LF spectral power values during the *Zen Garden* compared to control condition over 20-minute duration **e** Unresponsive participants' averaged LF spectral power also increased during the *Zen Garden* condition compared the control condition (p=0.009, *W*=63, *r*=0.59, 95% CI [0.06, 0.86]).
*Zen Garden* and control cities conditions depicted in green and orange, respectively. Temporal plots (b,d) depict datapoints for each window size=1 minute and step size=5 seconds, averaged ± SEM. Removed breathing exercise epoch is highlighted in blue. For (c,e) each participant's data is represented by connecting blue points. The boxplot inside the violin plot corresponds to the interquartile range, the median is depicted with a black dot, the vertical green and orange lines correspond to the probability density function. N=27 all participants, n=16 responsive and n=11 unresponsive participants. Wilcoxon signed rank test, adjusted with False Discovery Rate correction.

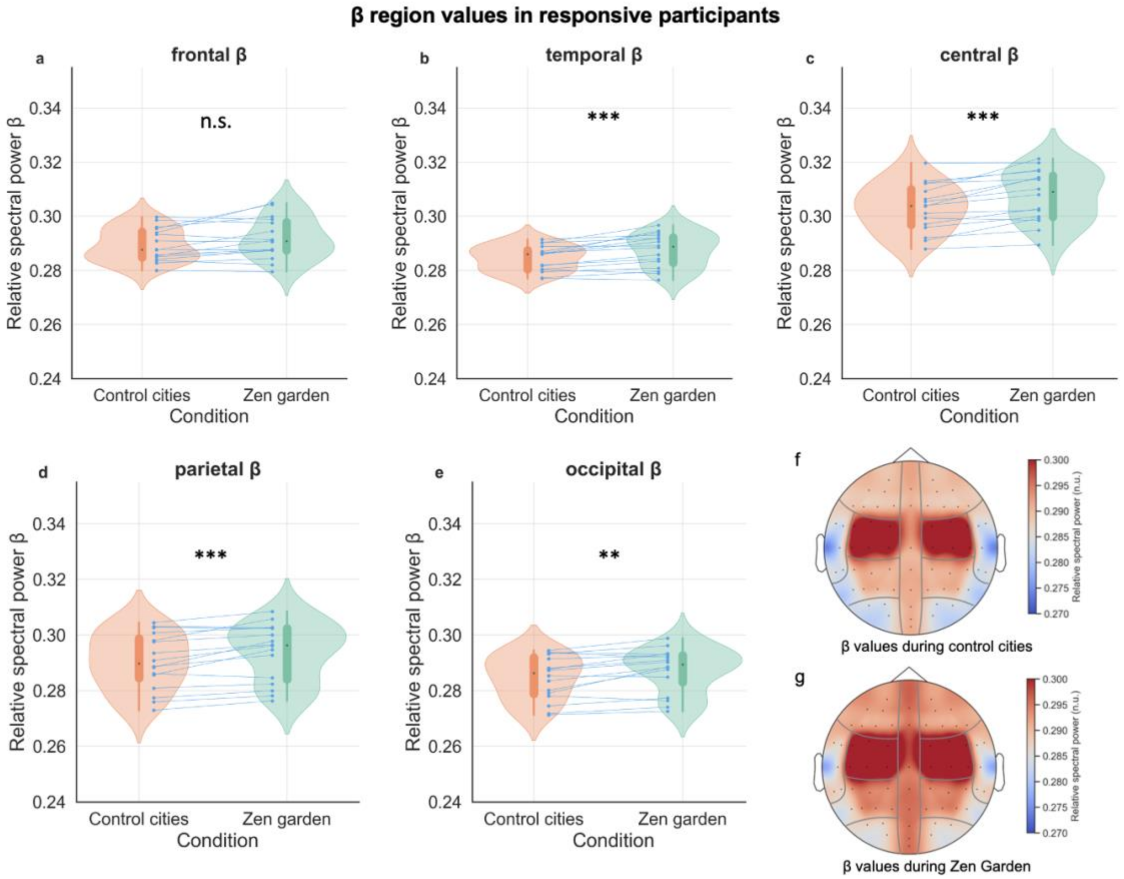


**Fig. S5. Conditions effects on beta regions in responsive participants. a,b,c,d,e** Beta relative spectral power (13-30 Hz) variations between *Zen Garden* and control video conditions in responsive participants in **a** frontal (p=0.07), **b** temporal (p=0.001, *W*=130, *r*=0.30, CI [-0.12, 0.64), **c** central (p=0.0009, W=134, *r*=0.28, CI [-0.14,0.62]), **d** parietal (p=0.001, *W*=130, *r*=0.17 [-0.24, 0.53]), **e** occipital (p=0.006, *W*=122, *r*=0.18, CI [-0.24, 0.54]) regions. Across conditions, temporal, central parietal and occipital regions displayed statistically significant difference in this responsive group, whereas the frontal region did not. **f,g** Topographic representation illustrating averaged beta relative spectral power in responsive participants **f** during the control condition and **g** during the Zen Garden condition. The groups of electrodes are averaged as in Fig. S3.
*Zen Garden* and control cities description conditions depicted in green and orange, respectively. Each participant's data is represented by connecting blue points (N=16 participants). The boxplot inside the violin plot corresponds to the interquartile range, the median is depicted with a black dot, the vertical green and orange lines correspond to the probability density function. Wilcoxon signed rank test, adjusted with False Discovery Rate correction.

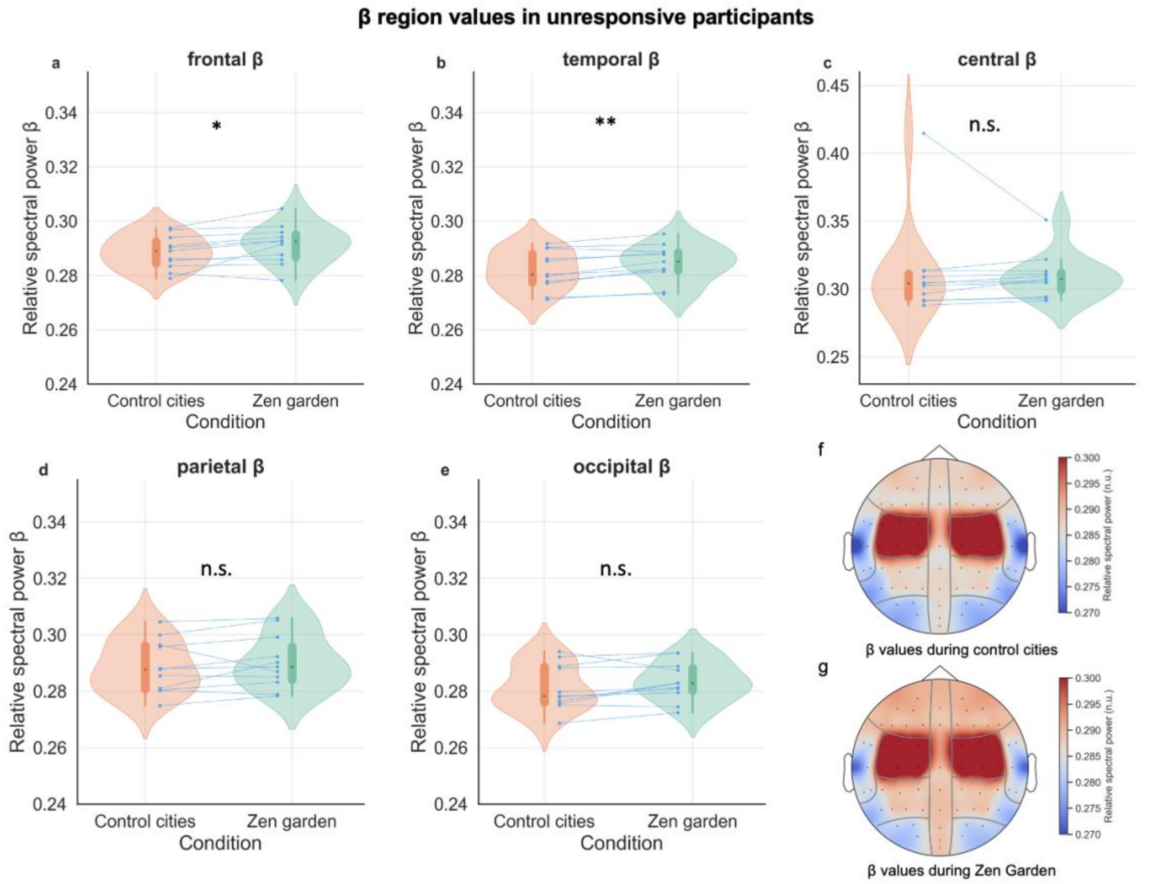


**Fig. S6. Conditions effects on beta regions in unresponsive participants. a,b,c,d,e** Beta relative spectral power (13-30 Hz) variations between *Zen Garden* and control video conditions in unresponsive participants in **a** frontal (p=0.04, *W*=58, *r*=0.26, CI [-0.26, 0.66]), **b** temporal (p=0.006, *W*=64, *r*=0.21 [-0.30, 0.62]), **c** central (p=0.07) , **d** parietal (p=0.14), **e** occipital (p=0.07) regions. Across conditions, the frontal and temporal regions displayed statistically significant difference in this unresponsive group. **f,g** Topographic representation illustrating averaged beta relative spectral power in unresponsive participants **f** during the control condition and **g** during the *Zen Garden* condition. The groups of electrodes are averaged as in Fig. S3.
*Zen Garden* and control cities description conditions depicted in green and orange, respectively. Each participant's data is represented by connecting blue points (N=11 participants). The boxplot inside the violin plot corresponds to the interquartile range, the median is depicted with a black dot, the vertical green and orange lines correspond to the probability density function. Wilcoxon signed rank test, adjusted with False Discovery Rate correction.

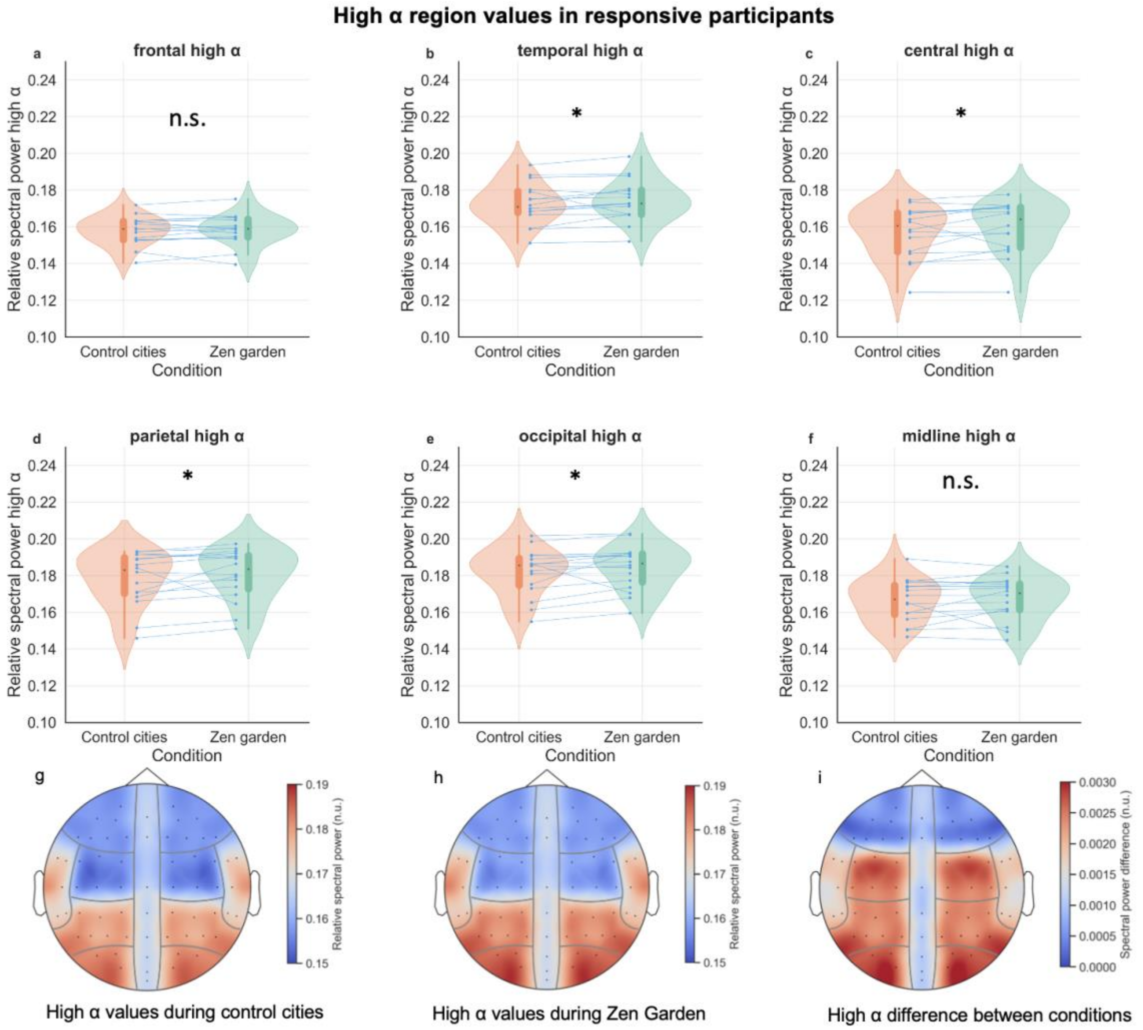


**Fig. S7. Supplementary Figure 7. Conditions effects on high alpha regions in responsive participants. a,b,c,d,e,f** High alpha relative spectral power (10-13 Hz) variations between *Zen Garden* and control video conditions in responsive participants in a frontal (p=0.49), **b** temporal (p=0.04, *W*=112, *r*=0.11, CI [-0.30, 0.48]), **c** central (p=0.04, *W*=110, *r*=0.16, CI [-0.29, 0.52]), **d** parietal (p=0.04, *W*=112, *r*=0.12, CI [-0.28,0.49]), **e** occipital (p=0.04, *W*=116, *r*=0.16, CI [-0.26,0.52]) and **f** midline (p=0.30) regions. Across conditions, temporal, central, parietal and occipital regions displayed statistically significant difference in this responsive group, whereas the frontal region did not. **g,h,i** Topographic representation illustrating averaged high alpha relative spectral power in responsive participants **g** during the control condition and **h** during the *Zen Garden* condition, **i** contrast between conditions (*Zen Garden* – control conditions). The groups of electrodes are averaged as in Fig. S3.
*Zen Garden* and control cities description conditions depicted in green and orange, respectively. Each participant's data is represented by connecting blue points (N=16 participants). The boxplot inside the violin plot corresponds to the interquartile range, the median is depicted with a black dot, the vertical green and orange lines correspond to the probability density function. Wilcoxon signed rank test, adjusted with False Discovery Rate correction.

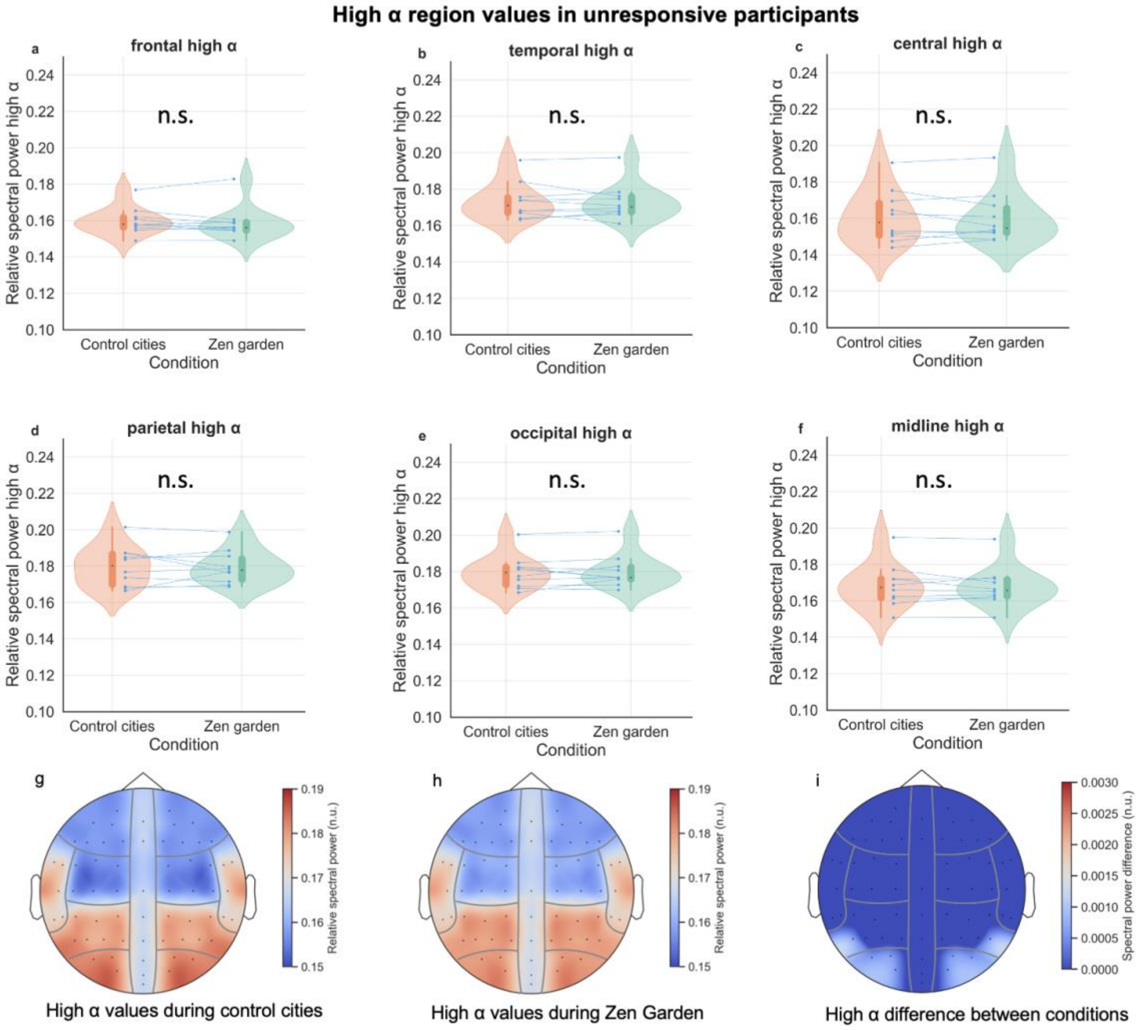


**Fig. S8. Conditions effects on high alpha regions in unresponsive participants. a,b,c,d,e,f** High alpha relative spectral power (10-13 Hz) variations between *Zen Garden* and control video conditions in unresponsive participants in **a** frontal (p=1), **b** temporal (p=1), **c** central (p=1), **d** parietal (p=1), **e** occipital (p=1) and **f** midline (p=1) regions. **g,h,i** Topographic representation illustrating averaged high alpha relative spectral power in unresponsive participants **g** during the control condition and **h** during the *Zen Garden* condition, **i** contrast between conditions (*Zen Garden* – control conditions). Wilcoxon signed rank test, adjusted with False Discovery Rate correction). The groups of electrodes are averaged as in Fig. S3.
*Zen Garden* and control cities description conditions depicted in green and orange, respectively. Each participant's data is represented by connecting blue points. N=11 participants, one participant was excluded after identifying outliers based on the 5%-95% range calculation, resulting in n=10 participants. The boxplot inside the violin plot corresponds to the interquartile range, the median is depicted with a black dot, the vertical green and orange lines correspond to the probability density function. Wilcoxon signed rank test, adjusted with False Discovery Rate correction.

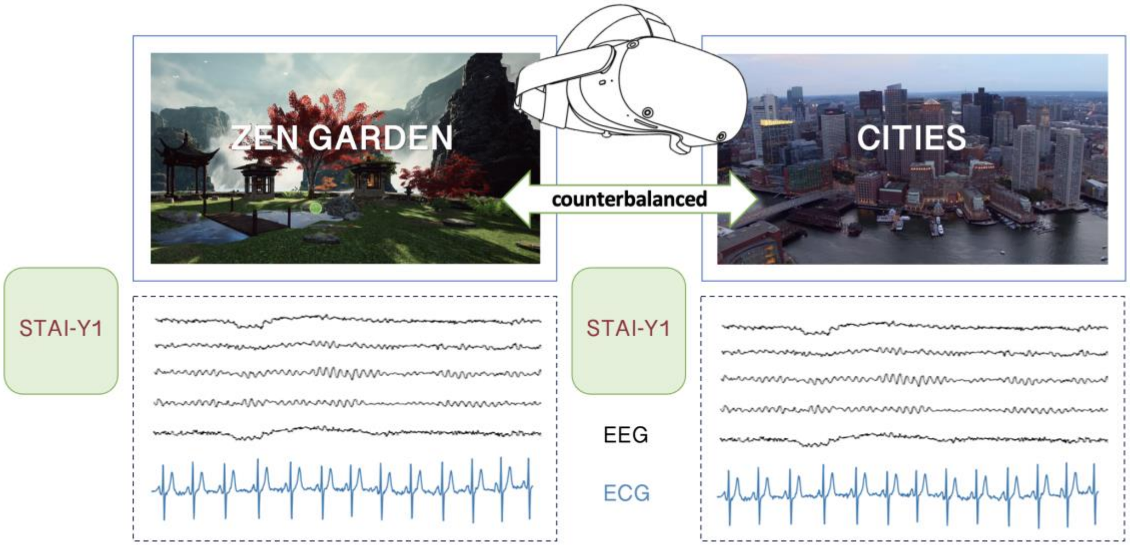


**Fig. S9. Experimental design.** Illustration of the two videos displayed in the virtual reality headset, simultaneously with EEG and ECG acquisition. Depiction of the STAI-Y1 questionnaires completed at baseline and after the *Zen Garden* condition.

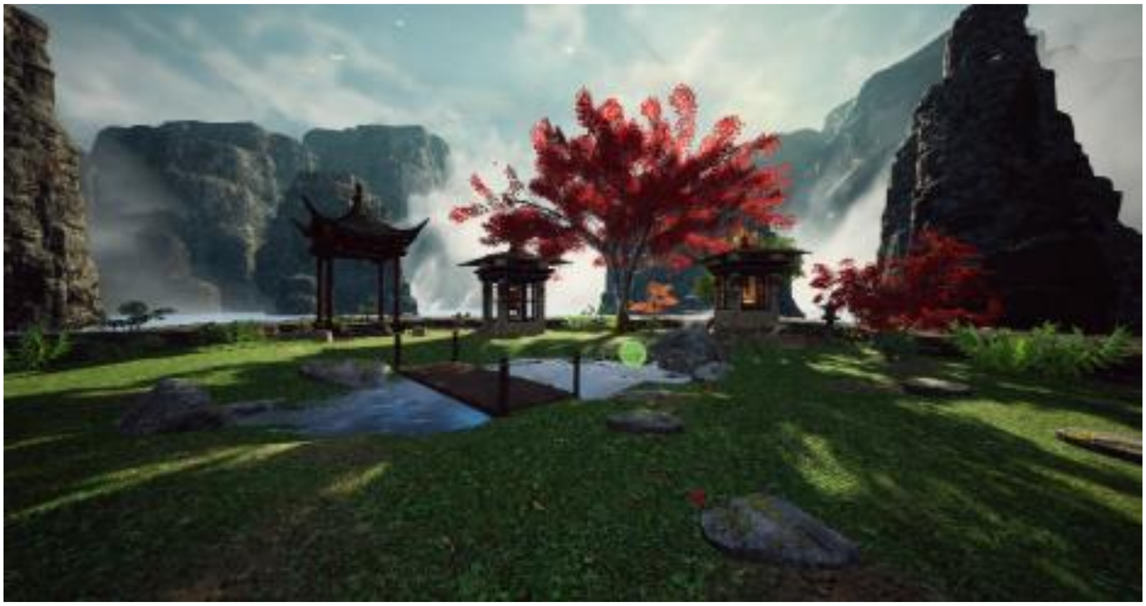

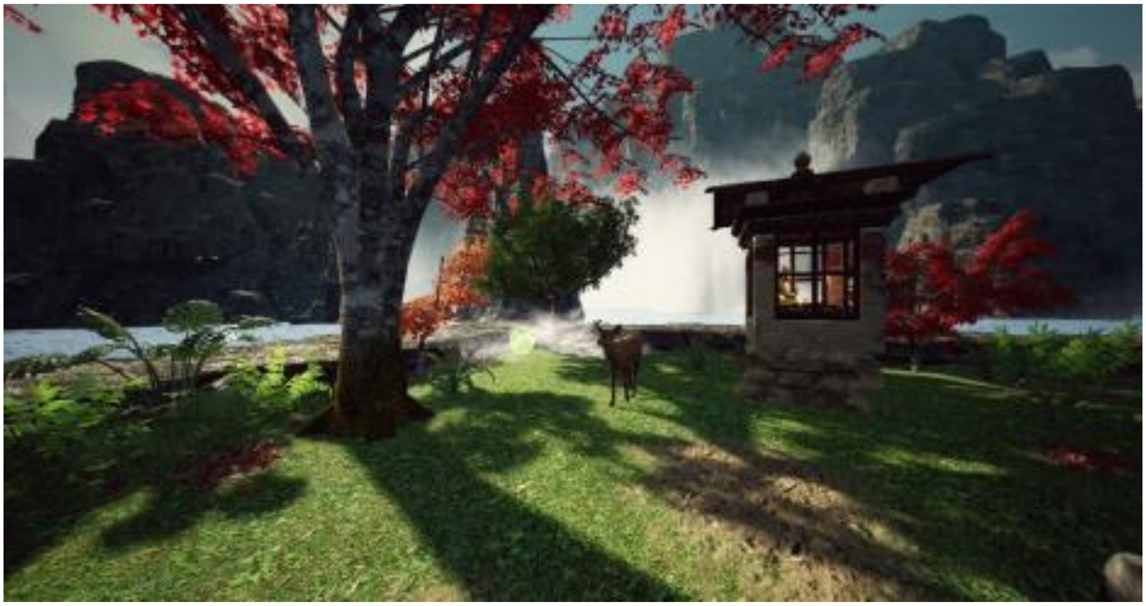

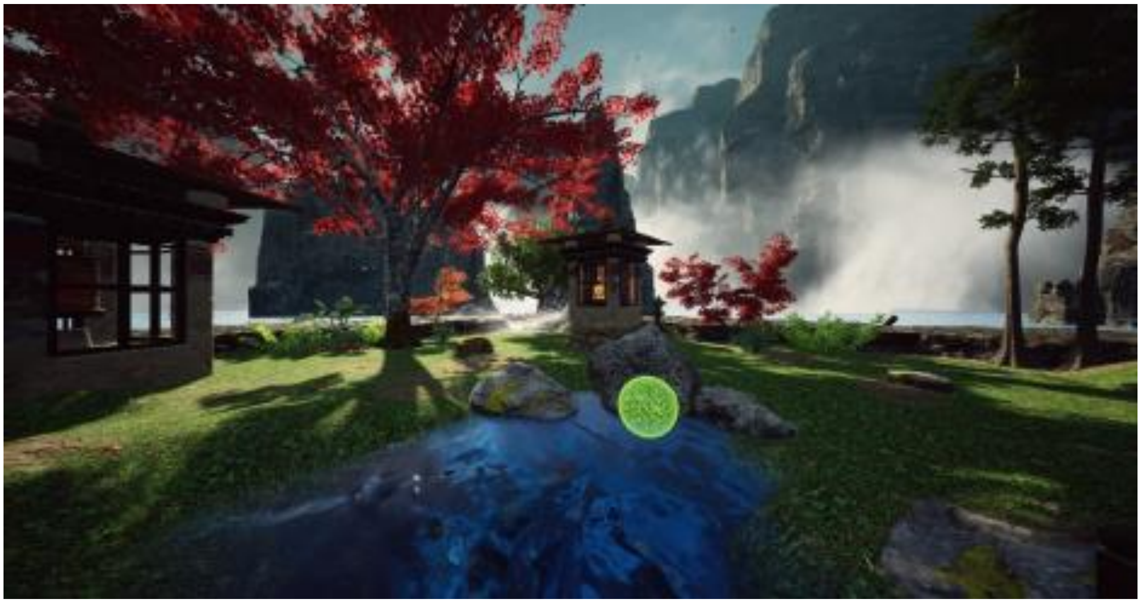

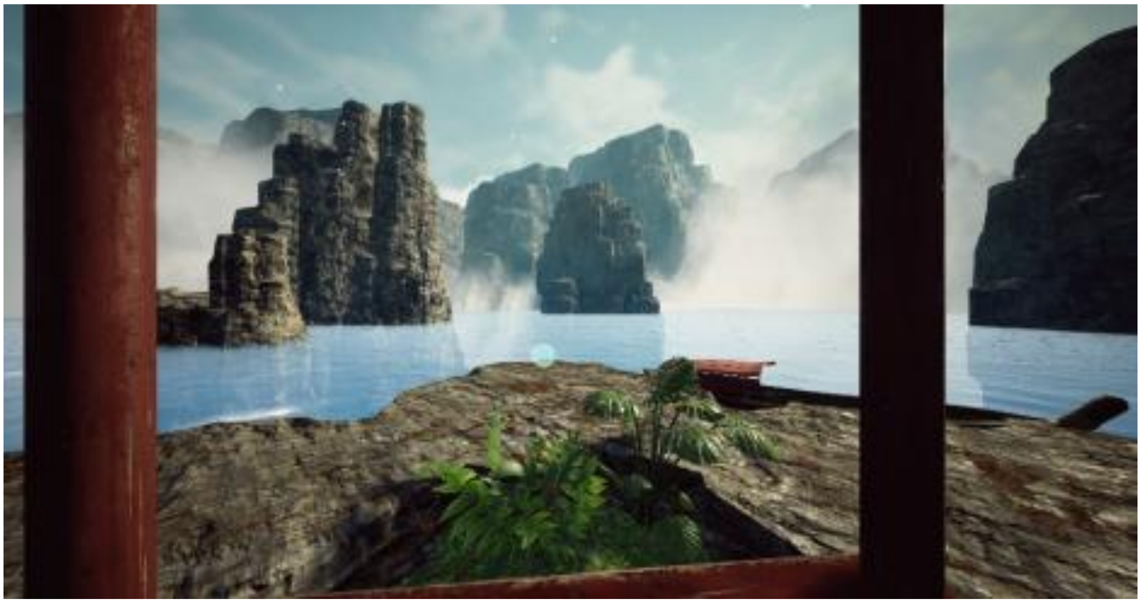

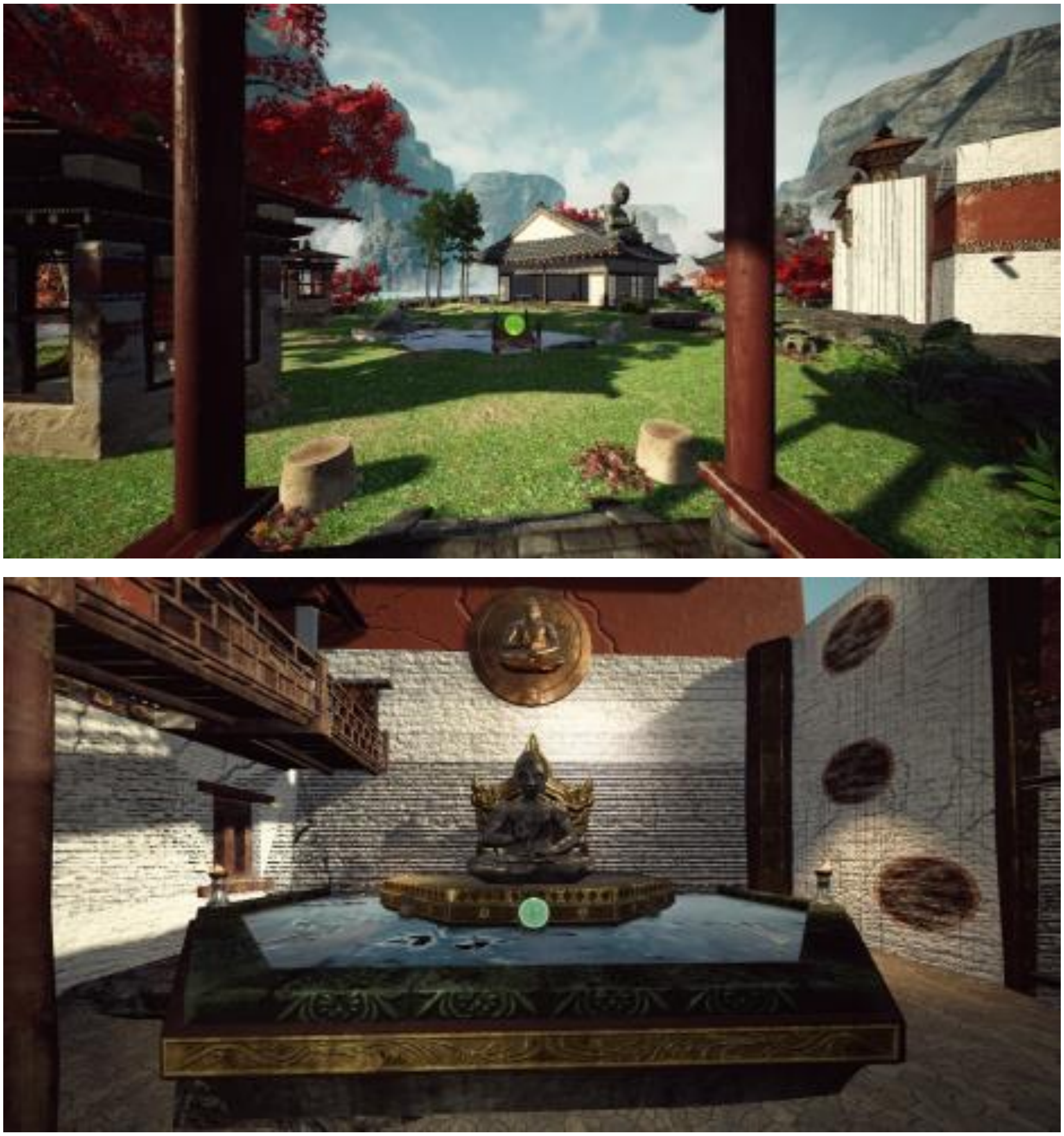

**Fig. S10.  Snapshots of the virtual *Zen Garden* environments.**

**Table S1.** Timestamps and scripts used in the *Zen Garden* virtual environment.

| Timestamp | Script |
|---|---|
| **00:02:40** | Let my voice guide you |
| **00:02:51** | Allow yourself to be lulled by the words, sounds, images, and all the pleasant sensations… |
| **00:03:02** | If your eyelids feel heavy, feel free to let your eyes close, and you can always return to this journey whenever you like |
| **00:03:32** | In the early morning, nature comes alive with energy |
| **00:03:53** | You can feel the warmth of the sun on your shoulders |
| **00:04:13** | The water sparkles and creates a gentle rhythm that matches your steps |
| **00:04:33** | You think to yourself: I'm moving forward with energy now |
| **00:05:05** | A harmonious interplay between the wind rustling through the leaves |
| **00:05:16** | The energy of the earth and the rock |
| **00:05:25** | The gentle ripples on the surface of the lake |
| **00:05:44** | You move forward slowly, in harmony with your surroundings |
| **00:06:04** | You feel increasingly calm |
| **00:06:22** | Your gaze is captivated |
| **00:06:34** | You feel as though you're part of a peaceful world |
| **00:07:02** | Back in the garden, a presence steps out of the shadows |
| **00:07:12** | A deer approaches |
| **00:07:31** | Calm and serene |
| **00:08:02** | You think: I feel safe |
| **00:08:30** | The bridge |
| **00:08:40** | The refreshing droplets of water on your face |
| **00:08:58** | You admire the sparkling spring as it flows and meanders through the garden |
| **00:09:12** | Among the ferns and rocks |
| **00:09:33** | Your gaze takes in the rock extending above the lake |
| **00:09:45** | You observe every detail of this beautiful landscape |
| **00:09:55** | The soft scent of water lilies fills the air |
| **00:10:15** | Breathe deeply |
| **00:10:43** | The temple is ready to welcome you |
| **00:11:03** | Clouds drift silently across the blue sky |
| **00:11:22** | Your steps take you from smooth rock to soft grass |
| **00:11:50** | Mist blends the lake and the sky |
| **00:12:09** | You continue confidently through the garden |
| **00:12:20** | You think: I walk and breathe with ease |
| **00:12:48** | You follow a path leading to the main temple |
| **00:12:59** | In the shade of pagodas, sheltered from the breeze |
| **00:13:18** | The area is lush with plants in shades of red and green |
| **00:13:48** | The rock shows tones of gray and brown, reflecting the dancing shadows of the trees |
| **00:14:03** | You walk towards the temple under the warm sun |

| | |
|---|---|
| **00:14:27** | You sit on a eucalyptus bench, surrounded by its soothing aroma |
| **00:14:49** | You take time to appreciate the harmony of the elements |
| **00:14:57** | The meeting of water, earth, and sky |
| **00:15:15** | You think: I savor all this beauty |
| **00:15:29** | Walking carefully along the path by the rocks |
| **00:15:43** | Your breath feels refreshed by the mist from the lake |
| **00:15:59** | Your feet return to the garden’s grass |
| **00:16:17** | Breathe, as the surface of the pool gently ripples |
| **00:16:38** | From the treetops, the silent flight of butterflies accompanies you |
| **00:17:19** | The path is easy |
| **00:17:30** | Your steps are light |
| **00:18:17** | The door to time opens, inviting you inside |
| **00:18:50** | Welcome |
| **00:19:04** | You can sit now and simply be |
| **00:19:39** | You’re surrounded by a deep sense of harmony |
| **00:19:49** | You feel calm and at peace |

**Dataset (separate file).** The excel file contains: Datasets S1 to S3 and S5 to S8, datasets Figure 1 to 6, datasets Table 1
